# Conformational Heterogeneity of RNA Stem-Loop Hairpins Bound to FUS RNA Recognition Motif with Disordered RGG Tail Revealed by Unbiased Molecular Dynamics Simulations

**DOI:** 10.1101/2022.08.16.504132

**Authors:** Pavlína Pokorná, Miroslav Krepl, Sébastien Campagne, Jiří Šponer

**Affiliations:** Institute of Biophysics of the Czech Academy of Sciences, Královopolská 135, 612 65 Brno, Czech Republic; National Centre for Biomolecular Research, Faculty of Science, Masaryk University, Kamenice 5, 625 00 Brno, Czech Republic; Inserm U1212, CNRS UMR5320, ARNA Laboratory, University of Bordeaux, 146 rue Léo Saignat, 33076 Bordeaux Cedex, France

## Abstract

RNA-protein complexes use diverse binding strategies, ranging from structurally well-defined interfaces to completely disordered regions. Experimental characterization of flexible segments is challenging and can be aided by atomistic molecular dynamics (MD) simulations. Here we used extended set of microsecond-scale MD trajectories (400 μs in total) to study two FUS-RNA constructs previously characterized by NMR spectroscopy. The FUS protein contains well-structured RNA Recognition Motif domain followed by presumably disordered RGG tail and bind RNA stem-loop hairpins. Our simulations provide several suggestions complementing the experiments but also reveal major methodological difficulties in studies of such complex RNA-protein interfaces. Despite efforts to stabilize the binding via system-specific force-field adjustments, we have observed progressive distortions of the RNA-protein interface inconsistent with experimental data, as in detail documented. We further propose that the dynamics is so rich that its converged description would not be achievable even upon stabilizing the system. Still, after careful analysis of the trajectories, we have made several suggestions regarding the binding. We identify substates in the RNA loops which can explain the NOE data. The RGG tail localized in the minor groove remains disordered, sampling countless transient interactions with the RNA. There are long-range couplings among the different elements contributing to the recognition, which can lead to allosteric communication throughout the system. Overall, the RNA-FUS systems form dynamical ensembles that cannot be fully represented by single static structures. Thus, albeit imperfect, MD simulations represent a viable tool to investigate them.

## Introduction

RNA molecules continuously interact with proteins. RNA-binding strategies employed by proteins range from using well-defined globular domains with canonical binding sites to disordered domains with dynamic or even fuzzy binding modes.^1–3^ Many RNA-binding proteins form large complexes composed of several functional modules/domains, which can cooperate in RNA binding. Coupling between order and disorder is fundamental for function of many of them.^4^

Molecular dynamics (MD) is an important tool to study dynamics in biomolecular systems. It provides unique data on biomolecular structural dynamics, folding and binding. Indeed, single-strand RNA binding to different globular RNA-binding protein domains was studied in many previous MD studies and is well-described by current versions of AMBER force field^5, 6^ while larger RNA-protein complexes are also being successfully studied.^7^ Yet, MD simulations often suffer from errors introduced by the empirical force fields as well as by the inaccessibility of large-scale sampling. Due to sampling limitations, potential inaccuracies in the starting structures may not have enough time to get corrected even with accurate force fields. Even description of small RNAs such as short single-strands and small RNA hairpins is still far from ideal with current state-of-the art force fields.^6, 8–10^ Another known issue of contemporary biomolecular force fields is their incapability to simultaneously describe folded and disordered protein regions.^11, 12^ Biomolecular complexes are, however, commonly composed of multiple different structural motifs and combine different degrees of disorder.^13^

Here, we report an MD simulation study of such a heterogeneous system, two complexes of the RRM-RGG domain of Fused in Sarcoma protein (**FUS**, also known as Translated in Liposarcoma, TLS, or hnRNP P2) bound to two different RNA hairpins (***Figure 1***). FUS is an RNA/DNA binding protein from the family of heterogeneous nuclear ribonucleoproteins (hnRNPs), which forms dynamic complexes with nascent RNA transcripts and controls their maturation, transport and translation while also being involved in DNA damage repair.^14, 15^ FUS mutations affect mRNA processing and typically result in FUS aggregation in cytoplasm which leads to severe neurodegenerative diseases such as amyotrophic lateral sclerosis, frontotemporal dementia, or essential tremor.^16–18^ FUS protein consists of a N-terminal low-complexity QGSY-rich prion-like domain followed by a glycine-rich region, three disordered RGG motifs encompassing two globular RNA-binding domains – an RNA Recognition Motif (RRM) domain and a Zinc Finger (ZnF), and a nuclear localization signal at C-terminus (***Figure 1***a). The RGG domains bind to RNA, DNA, and proteins and enable phase-separation, along with the prion-like domain.^19–23^ The two globular domains enable FUS to bind to RNA in a bipartite mode with context-dependent synergy.^24, 25^ While ZnF binds specifically to GGU motifs in a single stranded context, the RRM interacts with stem loops or hairpins with a poor sequence selectivity, indicating rather shape-specific recognition.^25^ The RRM binds RNA hairpins with YNY motif (Y=C,U; N=any nucleotide) at the 3’-end of the loop while some ssRNAs with the same motif can also be bound.^25–27^ The RGG extension at the RRM C-termini is not essential for RRM-RNA binding but increases RNA-binding affinity by 6-fold via enthalpy contribution though the affinity is still low (8-9 μM).^25, 26^ The solution structures of FUS RRM-RGG bound to two RNA hairpins (***Figure 1***b and d), namely the spliceosomal U1 snRNP stem-loop 3 (U1 SL3) hairpin and the hnRNPA2/B1 pre-mRNA hairpin (PDB IDs 6SNJ and 6GBM, respectively).^25, 26^ FUS RRM binds RNA using non-canonical RRM interface with just one aromatic stacking platform^25, 28^ while the major groove of RNA stem is contacted by the KK loop (loop α1-β2). The RGG2 extension binds dynamically into the RNA minor groove and can thus orient the ZnF to its binding site with respect to RRM.^26^ Both solution structures (PDB IDs 6SNJ and 6GBM) are currently the only experimentally resolved RNA-protein structures which show flexible binding of RGG. They contrast with the X-ray structures of FMRP RGG peptide bound to RNA quadruplex-duplex junction (PDB ID 5DEA)^29^ and of RGG extension of SF3A1 ubiquitin-like domain binding to rigid U1 snRNP stem loop 4 (PDB ID 7P0V)^30^ where the RGG segments are structured. In those two complexes, RGGs bind specifically to bases in the major groove of RNA helices. In the case of SF3A1, the RGG tail is required for RNA-protein binding while in the case of FUS, the tail constitutes an auxiliary element of the RNA binding interface.

**Figure 1:**
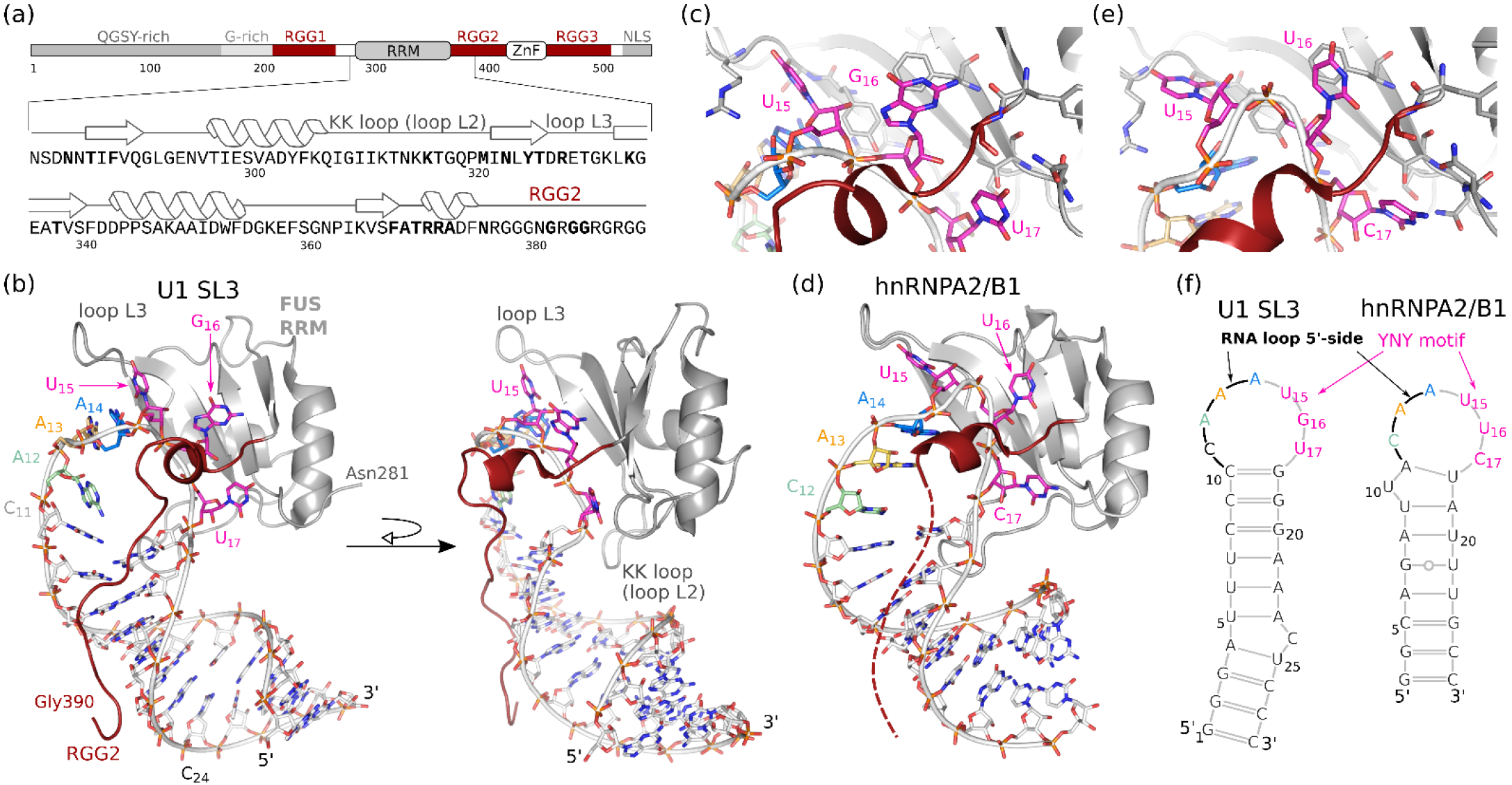
Overview of FUS-RNA binding. (a) Domain composition of the FUS protein and sequence of the construct used in this study. (b) Structure of the FUS complexes with U1 SL3 as resolved by NMR (PDB ID 6SNJ). (c) Detailed view on the YNY motif recognition in the FUS-U1 SL3 complex. (d) Solution structure of the FUS RRM in complex with hnRNPA2/B1 a (PDB ID 6GBM). The RGG extension was shorter in FUS-hnRNPA2/B1 complex and was modeled in MD simulations (indicated by dashed line). (e) Detailed view on the YNY motif recognition in FUS-hnRNPA2/B1 complex. (f) Scheme of the U1 SL3 and the hnRNPA2/B1 RNA hairpins.

Thus, the FUS RRM-RGG-RNA system combines the globular RRM domain with disordered RGG tail and RNA with larger loop connected to the RRM-bound nucleotides. Both RNA hairpins differ also in properties of their stems, with one closing the loop with three GC base pairs and the other one with a flexible stem (***Figure 1***f).

Our MD simulation study complements the existing NMR investigations and provides suggestions about the properties of the FUS-RNA complexes, such as coexistence of RNA substates which cannot be resolved in the ensemble-averaged experiments. At the same time, the simulations give unprecedented insights into uncertainties, problems and limitations of MD technique, which to our opinion are rarely documented in contemporary literature.^31^ The FUS systems turned out to be very challenging for MD simulations. In fact, despite that we have accumulated more than 400 μs of simulation data, the overall sampling of our simulation dataset remains in an extremely data-poor regime. The force-field description suffers from visible limitations resulting in progressive deterioration of the simulated structures with no sign of convergence while there are also genuine uncertainties in the experimental starting structures. Limited improvement was achieved by employing several system-specific force-field adjustments. In addition, simulations of the two different RNA hairpins, U1 SL3 and hnRNPA2/B1, show different structural stabilities of the simulated interfaces. It illustrates the known fact that simulation performance for RNA-protein complexes cannot be generalized even for similar RNA-protein systems.^6, 31^ Still, the simulation dataset contains useful information, especially in those initial parts of simulations preceding onset of a significant corruption of the structures. While the imperfect force-field performance and limited amount of experimental data do not allow us to confidently draw the real structural ensemble of the complex, we believe that we provide plausible suggestions about its properties based on the sub-states revealed by MD simulations. We show that a single conformation does not reproduce all the experimental NMR data and the RNA loops sample multiple conformational sub-states. We document sub-states that explain NOE signals which are problematic to reproduce in MD ensembles. We also propose that the complex can be divided into different parts that have different dynamics and differently contribute to the binding. The RGG tail remains disordered when bound to the RNA so that the RGG-RNA bound state consists of extremely diverse set of fluctuating interactions. Nevertheless, the RGG tail remains broadly localized in the RNA minor grove.

## Methods

### Starting structures

We simulated the FUS RRM-RGG complex with two RNA stem loops: U1 stem loop 3 (SL3, PDB ID 6SNJ, human)^26^ and hnRNPA2/B1 stem loop (PDB ID 6GBM, human, **Figure 1**).^25^ RNA residue numbering used in this work was chosen such that residues of both hairpins that bind to the same RRM binding sites have the same numbers (e.g. G_16_ and U_16_, ***Figure 1***f). G_16_ of U1 SL3 corresponds to G106 of human U1 and U_16_ of hnRNPA2/B1 to U3419 of human hnRNPA2/B1 transcript variant B1. To distinguish RNA and protein residues we use single-letter codes with a subscript for nucleotides and three-letter codes for amino acids. The first model of NMR structures was used as a starting structure. All NMR models are very similar and the choice of the model typically does not affect simulations since the variations in the NMR ensemble are smaller than fluctuations in MD.^5^ FUS residues 281-390 were used in simulations, corresponding to RRM and ~20 amino acids of the RGG tail (***Figure 1***a). The RGG extension in the FUS-hnRNPA2/B1 complex was modelled accordingly to FUS-U1 SL3 complex and 400 ns simulation with the RRM+RNA part restrained was run to allow the RGG to accommodate. We have also used some of the stable structures found by MD simulations as starting structures for additional MD runs; see Supporting Information for further details and PDBs of the starting structures. Details on supplementary RNA-RGG simulations of FMRP and SF3A1 complexes can be found in Supporting Information.

### Used force fields

RNA was described by the standard AMBER OL3 force field.^32–35^ Concerning the protein part, majority of simulations is based on AMBER ff14SB protein force field^32, 33, 35, 36^ with cufix correction.^37, 38^ Cufix is a pair-specific adjustment of Lennard-Jones parameters (NBfix) for pairs of nitrogen and oxygen atoms of charged groups and for interactions of aliphatic carbons which decreases excessive solute aggregation and charge-charge interactions. We used the SPC/E water model^39^ and Li&Merz ions (12-6 hydration free energy set).^40^ We have also run some simulations without the cufix or with AMBER ff19SB protein force field^41^ and OPC water model^42^ with OPC-adapted Li&Merz ions. 2 kcal/mol HBfix^43, 44^ (see below) was used for moderate stabilization of H-bonds of two terminal base pairs of the RNA stems since they otherwise have tendency to unpair, fray and subsequently stick to the protein. Further, we have applied some other system-specific force-field adjustments as described below.

### Simulation setup

Simulations were performed with Amber18 and Amber20 packages.^45, 46^ All systems were equilibrated using standard protocols with gradual heating and decrement of heavy-atom restraints followed by an extended equilibration protocol with a 100 ns simulations with gradual decrement of all NOE NMR restraints for smoother relaxation of the restrained structure.^31, 47^ Production runs (3-10 μs long, multiple independent trajectories per system) were run with *pmemd.cuda* module.^48, 49^ The Langevin thermostat with a collision frequency of 5 ps^-1^ and Monte-Carlo barostat^50^ were used to maintain a temperature of 300 K and pressure of 1 bar, respectively, while SHAKE algorithm^51^ and hydrogen mass repartition scheme^52^ were applied, allowing a 4 fs time step. Velocities were randomized each 1 ns.

### Simulations sets and simulation ensembles used in this study

#### Preliminary simulation set

We have initially run simulations without any or just with subtle modifications of the system and force field (*preliminary* simulation set). They were done on timescales 0.5-5 μs with total time of ~140 μs for both systems combined. All preliminary FUS-hnRNPA2/B1 simulations were done without the modeled RGG extension. The FUS-U1 SL3 the preliminary simulation set includes also some simulations with cufix correction and simulations with ff19SB protein force field and OPC water model. Trajectories from the preliminary simulation set, however, were notably unstable. We have observed mainly excessive sticking of RNA stem to FUS RRM and/or pronounced progressive changes in the RNA loop and RNA-protein interface (unbinding of the YNY motif or formation of long-lived states with massive NOE violations). The unstructured N-terminal extension preceding Asn281 from FUS-U1 SL3 complex always detached from the domain and bound to the RNA stem with no sign of reversibility, which contradicts protein-protein NOE data between the N-terminal extension and the RRM (~60 NOE pairs). Binding of N-terminal extension to RNA also affects RNA structure and its position. Another issue for the FUS-U1 SL3 system was irreversible Arg383 binding on top of the RNA stem. We consider this Arg383 stacking interaction to be over-stabilized by the force field as dominant population of Arg383 stacking would lead to multiple NOE violations involving C_11_ (see Supporting Information for details).

#### System-specific adaptations of standard protocols which were used in the main set of simulations

Based on the *preliminary* simulation set, we have system-specifically tuned our simulations. We call simulations with these adjustments as the *main* set of simulations and the cumulative times of the *main* sets were 124 and 135 μs for U1 SL3 and hnRNPA2/B1 FUS complexes, respectively. Besides using cufix, we have used 2 kcal/mol HBfix^43, 44^ and 1kcal/mol NOEfix^53^ potentials to support specific interactions, namely two U_15_-protein H-bonds in FUS-U1 SL3 complex and some NOE pairs, as listed in Table 1. We have used six and seven NOEfixes for U1 SL3 and hnRNPA2/B1 FUS complexes, respectively, as the respective distances were clearly violated in all or most of the *preliminary* trajectories. Both HBfix and NOEfix are based on the same principle; they represent usually small-to moderate potential energy bias with forces acting only within selected distances (2 - 3 Å for acceptor-H H-bond distance for HBfix and *r - 1.3·r* for NOEfix, where *r* is the NOE upper-bound). Thus they tune but not restrain the interaction (see Supporting Information, Figure S1 for details). Note that we refrain from using NMR NOE restraints in the simulations as the respective distances do not always have to be below the restraint threshold in reality. Applying hard restraints in such cases leads to trapping the structure in only one conformation. The N-terminus from FUS-U1 SL3 complex was cut in both complexes and the Asn281(ND)-Pro344(O) H-bond was restrained to keep the remaining part of the N-terminus positioned on RRM. For the FUS-U1 SL3 simulations, we enhanced Arg383 sidechain-water interactions by scaling Lennard-Jones R_min_ for N-O_w_ atom pairs by 0.95 to prevent irreversible Arg383 binding on top of the RNA stem. All those adjustments helped to reduce the severe distortions in the trajectories though even after applying them we continued to observe slower interface disruptions (see the Results section).

**Table 1.**
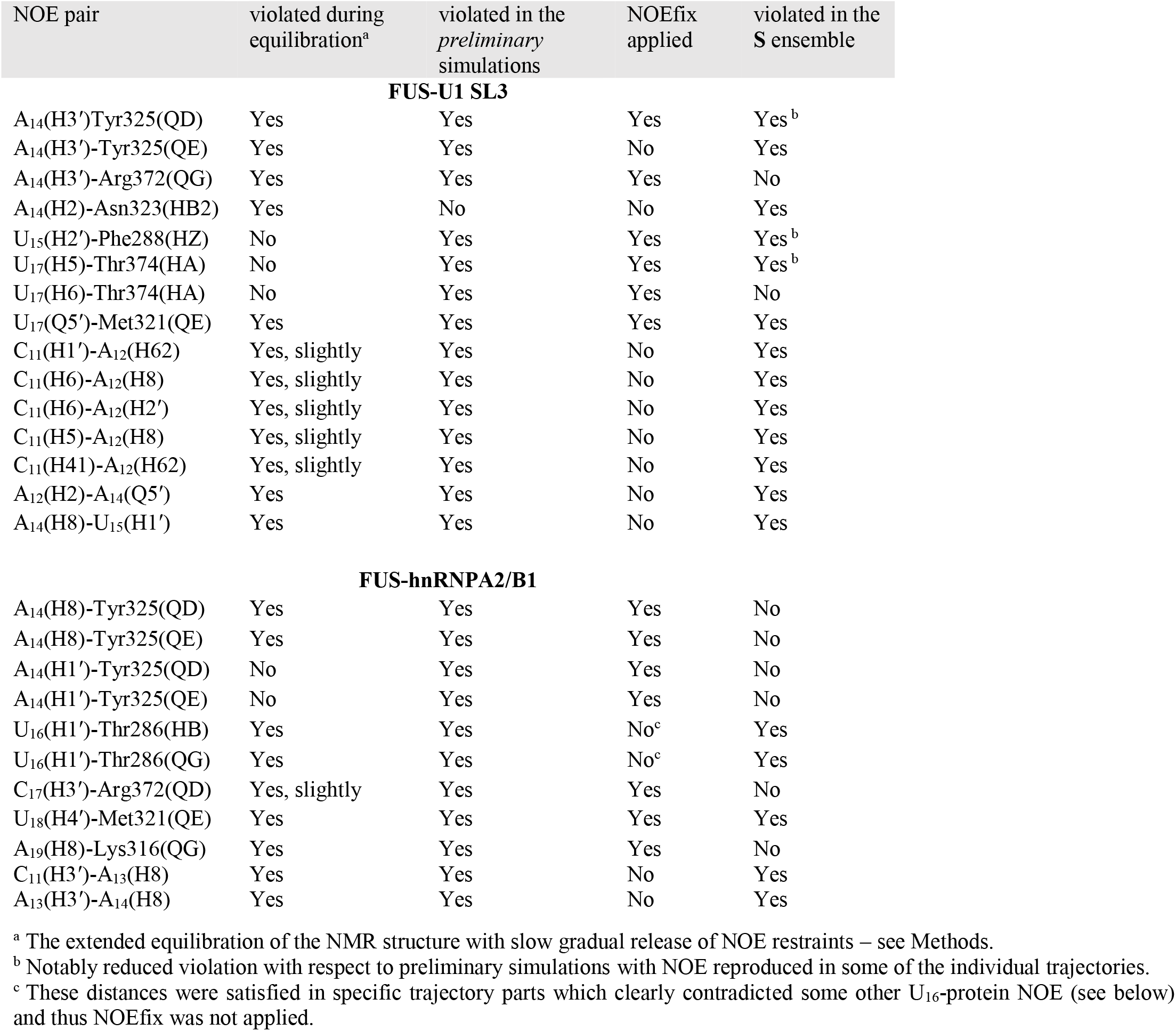
Overview of NOEs at the RNA-Protein Interface and in the RNA Loop’s 5’-end that Were Violated (> 1.0 Å) in the *Preliminary* Simulations and/or in the S Ensembles.

#### Selection of final simulation ensembles

Because of the observed disruptions we have divided trajectory parts from the *main* set of simulations into two ensembles, **S** and **U** (stable and unstable). The split of the trajectories into the **S** and **U** ensembles was done based on manual monitoring of the trajectories, distances and NOE violations. As the main criteria to classify a given trajectory portion to belong to the ensemble **S** we used the following requirements. For both systems we require that all nucleotides of the YNY motif remain bound in their pockets (including moderate fluctuations), KK-loop is in contact with RNA stem and structure of the loop 5’-end agrees with most of the NOEs (some violations are tolerated due to dynamical nature of the system). In addition, for the FUS-U1 SL3 system we require that Arg383 is not stacked on top of the RNA stem. In turn, for the FUS-hnRNPA2/B1 system we require no progressive changes in the stem structure, although fluctuations and temporary changes in the base pairing pattern are considered as unproblematic. We require a trajectory to be “stable” for at least 1 μs to include it in the **S** ensemble. Three trajectory parts from the FUS-U1 SL3 preliminary simulation set, that were stable, were also added to FUS-U1 SL3 **S** ensemble (Supporting Information Table S1). The final **S** ensembles contain 40.4 μs data for the FUS-U1 SL3 and 83.5 μs for the FUS-hnRNPA2/B1 (see ***Figure 2*** and Supporting Information Table S1).

**Figure 2:**
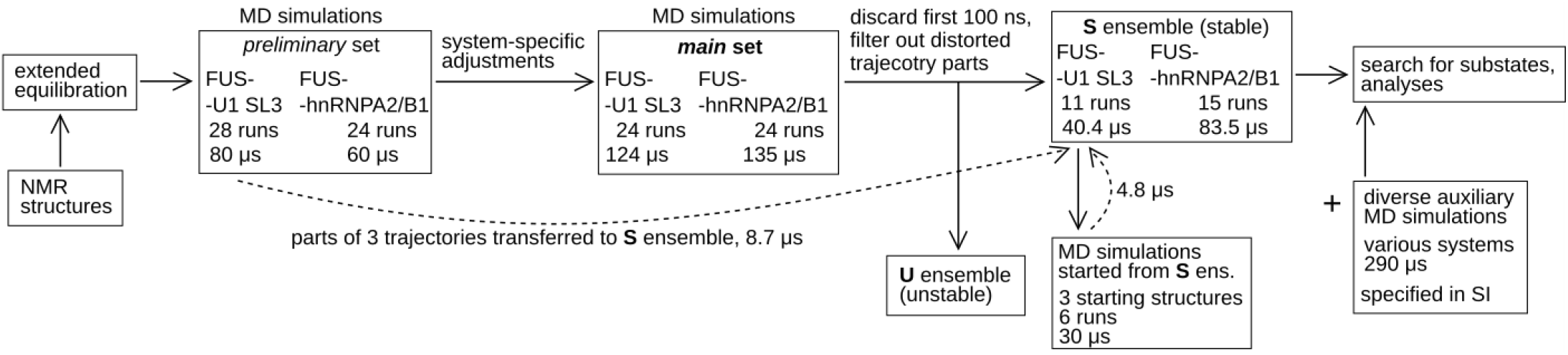
Overall scheme of all simulations and construction of the S ensembles used for the productive analyses. Individual simulations were typically run for 3-5 μs while the time-scale numbers shown in the boxes summarize the respective aggregate simulation time. Detailed list of all simulations is provided in Supporting Information, Table S1.

### Analyses

First 100 ns of all trajectories were discarded to reduce starting-structure bias. Simulation trajectories were analyzed using *cpptraj* module of Amber^54^ and VMD^55^. Figures were generated with PyMOL.^56^

NOEs were considered violated if the distance was > 1.0 Å above the experimental upperbound but we have also monitored violations > 0.3 Å. Besides the overall NOE violations in the ensemble and in individual simulations we have also analyzed instantaneous violations along the trajectories. Values of NOE violations in small trajectory parts are not representative for the overall agreement with experimental data but allow us to detect coupling of different NOEs and rare states. Also, simple NOE analysis over a whole trajectory can obscure structural distortions that occur later in the course of the trajectory since the average NOE signals will be biased by the near-starting structure values.

Since the overall simulation ensembles were not capable to satisfy many NOEs, we decided to search for rare simulation substates that could explain the observed signals, assuming that the simulations capture them properly but underestimate their population. To identify sub-states that could explain the significantly violated NOEs we have searched for snapshots where the respective contacts were formed and subsequently we analyzed the NOE signal smoothed with a window of 1 ns or 10 ps (for trajectory details) to include fluctuations around each snapshot. This allowed us to search for continuous trajectory portions where the signal was reproduced. We suggest that to estimate the minimal populations of the sub-state required to reproduce the experimental signals, including continuous trajectory portions is more appropriate than using just isolated snapshots.

As a check, we have also reweighted **S** ensembles with maximum entropy (ME) method.^57, 58^ This is an automatic procedure that already implicitly reflects coupling of NOEs. We could not use ME rigorously for obtaining a proper reweighted ensemble as we do not have sufficient amount of primary data and input ensembles of sufficient quality for the method (see Results and Supporting Information). However, we executed the reweighting procedure to obtain a qualified guess of which parts of trajectories could be important. For each violated NOE signal, we have then analyzed snapshots that contributed the most to the signal in the ME reweighted ensemble. Importantly, both ME and manual search identified the same sub-states, proving robustness of the substate analyses. We show an example of the analysis in the Supporting Information, Figure S6.

For the FUS-hnRNPA2/B1 complex a system with shorter RGG tail than for FUS-U1 SL3 was used for NOE data acquirement in the experiment. For simulations, we have extended the RGG tail to be of the same length in both systems. Thus, we did not consider NOEs reported for C-terminal residue in the experiment (Arg377) in our NOE analyses since its conformation is altered by the RGG extension. Additionally, we also observed NOE signals of preceding Arg372 to U_15_ to depend on the RGG tail length since these NOEs are violated in our simulation with extended RGG tail but not in *preliminary* simulations where the RGG length in FUS-hnRNPA2/B1 system was the same as in the experiment.

## Results & Discussion

### Simulations of FUS systems are in general unstable and trajectories divergent

Executing stable simulations of the FUS-RNA system turned out to be challenging. Based on the *preliminary* simulations, we made some adaptations of the basic simulation protocol for the *main* set of simulations. First, we have chosen the OL3+ff14SB+cufix+SPC/E combination of parameters as the basic force field (see Supporting Information for detail of force field selection). Then we have supported several RNA-protein H-bonds and NOE contacts at the interface using the HBfix/NOEfix approaches. Further, in FUS-U1 SL3 simulations, we have enhanced Arg383-water interactions to prevent the arginine from its clearly excessive stacking with the RNA stem (see Methods).

Despite these efforts, we have still observed major irreversible progressive changes at the RNA-protein interface and in the RNA loops in many trajectories (***Figure 3*** and Supporting Information, Table S3). Thus, we split the available trajectory portions into two subsets: subset **S** (stable, 40.4 μs for the FUS-U1 SL3 and 83.5 μs for the FUS-hnRNPA2/B1, ***Figure 2***) includes initial trajectory portions or full trajectories where the system can be considered to remain in a reasonable agreement with the experimental data (see Methods). Subset **U** (unstable) includes trajectory portions where the system shows already too large deviations. The differentiation between **S** and **U** was based on careful visual monitoring of all trajectories and analysis of many distances. Hallmarks of corrupted parts of trajectories in subset **U** are unbinding of the YNY-motif nucleotides from the RRM binding pockets, progressive rearrangements of the RNA loop 5’-end leading to states with massive NOEs violations (some NOE violations are tolerated since the simulations suggest coexistence of different states of the loop with conflicting NOEs, see below) and sticking of the RNA stem to RRM α2 helix or even to the U_17_/C_17_ nucleotides. Specifically in the FUS-U1 SL3 system, a visible artifact is a stiff continuous stacking of C_11_-U_15_ segment nucleotides at the loop 5’-end.

**Figure 3:**
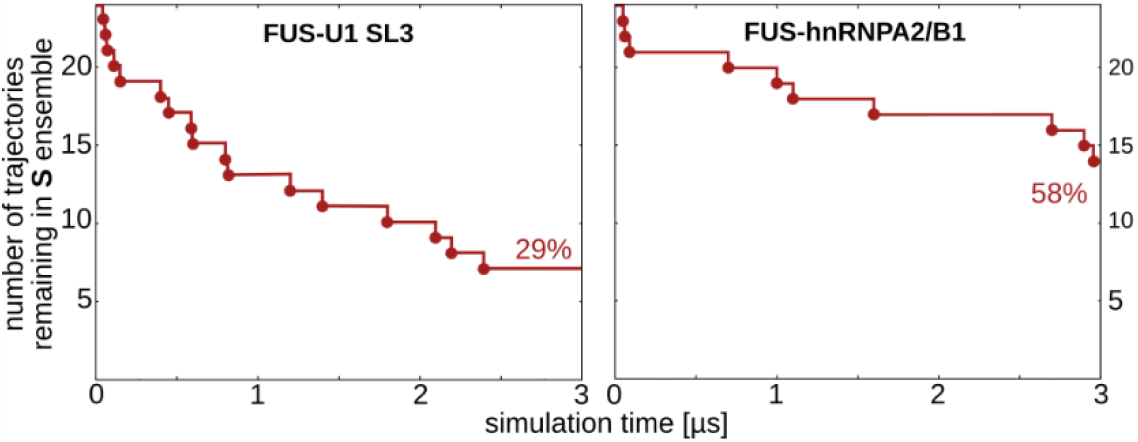
Cumulative graphs of the S to U transitions in the *main* set of simulations for all trajectories initiated from equilibrated NMR structures. For the FUS-U1 SL3 complex, the seven simulations that remained in the **S** ensemble till 3 μs were prolonged to 5 μs; one of them transited to the **U** ensemble. For the FUS-hnRNPA2/B1, fourteen simulations were prolonged and all of them remained classified as the **S** ensemble. This illustrates the more stable behavior of the FUS-hnRNPA2/B1 simulation set and the known fact that upon prolonging the simulations the probability of transitions becomes progressively reduced, as the simulated systems are often settling in longer-living substates. Structures from simulations that remained formally classified in the **S** ensemble at the end differ in many structural details, illustrating the insurmountable sampling limitations (Supporting Information Table S4 and Figure S7).

Generally, we did not observe any cases of coming back to subset **S** after reaching subset **U**, despite some occasional local structural improvements, accompanied, however, by distortions elsewhere. Prolongation of some corrupted trajectories to 10 μs revealed further and further departures from the starting structure as the errors continue to accumulate and propagate during the simulations. In addition, in simulations of both systems, we observed strong signs of autocorrelation,^59^ i.e., different long-lived states were sampled in different trajectories depending on the early trajectory developments (Supporting Information Table S4 and Supporting Information Figure S7). Occasionally, we observed transitions to structures sharing qualitative similarities of some monitored features with other trajectories. However, these trajectories were still sampling different overall conformations due to differences in other parts of the structures (e.g. RGG tail conformation). The system is beyond the convergent sampling capabilities of contemporary computers.

Although some simulations (of variable length) were still free of major distortions at their ends, we suggest that the force field is not able to provide a balanced overall description of the FUS-RNA complexes. This precluded substantial extensions of the trajectories or use of enhanced sampling methods. Thus, our strategy was the following. We based our productive analyses exclusively on the **S** ensembles, from which we identified structures that could contribute to the real conformational space of the FUS-RNA complexes. We assume that the initial stages of simulations represent a mix of real dynamics and force-field artifacts. We assume that the **S** ensembles may provide some important insights into the system, especially in view of the genuine uncertainty of the experimental data. In view of the overall simulation behavior, we refrained from using common methods (such as Principal Component Analysis or Markov State Modelling) which could mask the natural limitations of the simulations and are to our opinion not justified with the present systems. While the approach may seem excessively cautious (compared to practices sometimes used in contemporary RNA-protein MD literature), we aimed to avoid over-interpretations while seeking for options to use MD simulations even for RNA-protein systems whose description is difficult. We openly describe the difficulties as we believe it is an important part of research.

### FUS-hnRNPA2/B1 simulations are more stable than FUS-U1 SL3 simulations

The above-mentioned distortions were notably more pronounced in the FUS-U1 SL3 complex compared to the FUS-hnRNPA2/B1 (*Figure 3*). The different simulation stabilities of the two complexes could be a combination of three factors which are difficult to differentiate: existence of a real coupling between properties of the stem and loop length, strains in the ensemble-averaged starting structures, and imperfect force-field description of the stem-loop junction. The U1 SL3 stem has three rigid GC WC base pairs closing the loop while hnRNPA2/B1 stem has a dynamic U_10_/A_11_/U_18_ element followed by two AU WC base pairs and the GU wobble (***Figure 1***f). Simulations of isolated as well as FUS-bound hnRNPA2/B1 hairpin show reversible remodeling of the base pairing up to GU wobble (see Supporting Information for details). The better adaptability of the hnRNPA2/B1 stem could be one of the ways how this system compensates for strains in the starting structure and for force-field imbalances resulting in the overall better simulation behavior. The U1 SL3 stem with three rigid CG base pairs may be less capable of buffer tensions in the adjacent loop region. Indeed, when we mutated the three U1 SL3 GC base pairs into isosteric but more flexible AU base pairs, we observed strong buckling of the uppermost loop-closing AU base pair in simulations, suggesting some strain in the starting structure at the stem-loop junction. When one of the loop bases bulged out, the AU buckling was eliminated. Bulging out of the bases can thus also be a mechanism for U1 SL3 to release the strain that might be present in the starting structures. More details are in Supporting Information.

### Heterogeneous dynamical ensembles can reproduce the NMR data

As explained on particular examples below, we assume that the studied systems are dynamical and cannot be represented by just one structure satisfying all NOEs simultaneously. The potential extent of the real dynamics is documented by ***Figure 4***, which shows examples of time development of five RNA-protein inter-hydrogen distances in the FUS-U1 SL3 **S** ensemble. For ***Figure 4***, we have selected distances which show large fluctuations but still satisfy the NOE derived distances in the **S** ensemble. Some of the distances are above their upper bound limits in the majority of the trajectory parts and snapshots. On the right side of ***Figure 4***, we show the range of these distances in the NMR-refined ensemble, which is strikingly smaller compared to the distance ranges sampled in the simulations. Obviously, the extent of dynamics in simulation data can be biased by (and excessive due to) force-field approximations and imbalances. Still, the data clearly show that, in principle, the NOE signals for the studied system could be produced by very heterogeneous and dynamical ensembles. The more diverse the real ensemble is, the more likely the ensemble averaging introduces strains into the static predicted structures satisfying all the primary experimental data.

**Figure 4:**
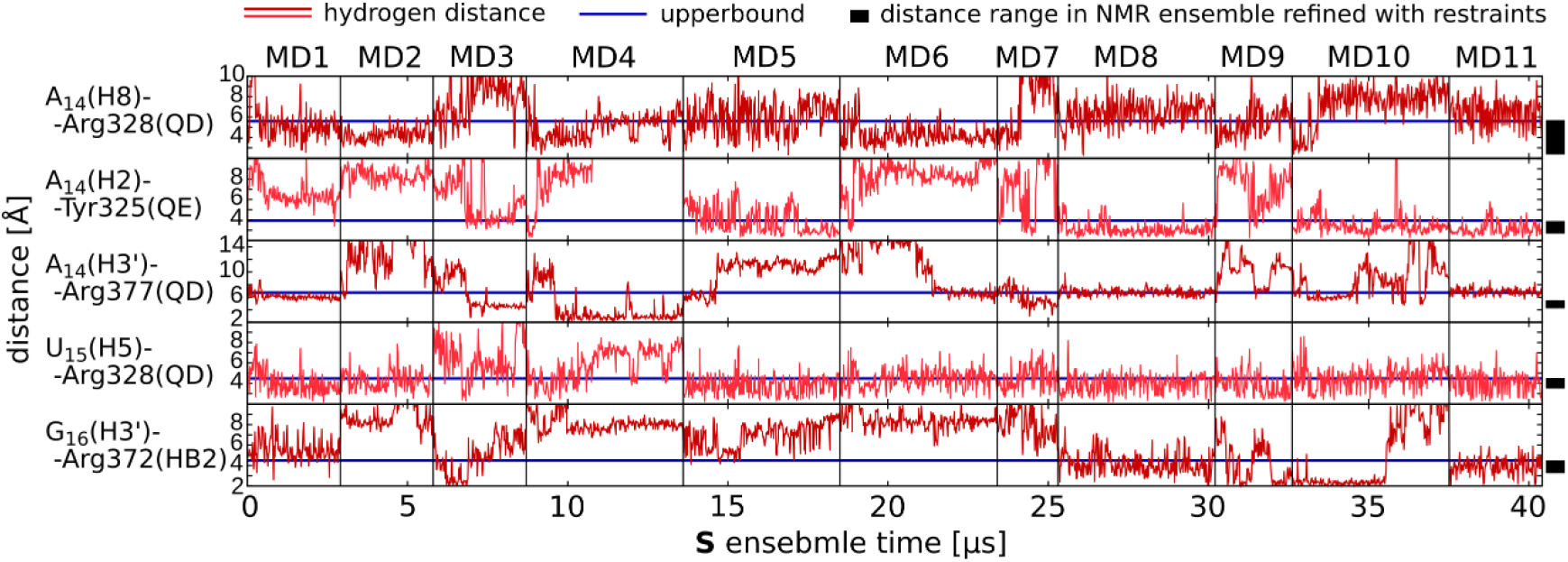
Time development (eleven simulations forming the complete S-ensemble, separated by the vertical lines) of selected FUS-U1 SL3 RNA-protein inter-hydrogen distances for which the NOE signals were measured. For hydrogen groups (e.g., QD), r^-6^ average of the respective hydrogens (e.g. HD2 and HD3) is plotted. The ranges of distances in the NMR-refined structural ensemble^24^ are shown as the black rectangles on the right side of the figure. For this graph, we selected distances that are substantially fluctuating while not being violated in the **S** ensemble.

Thus, we did not use any standard NOE restraints in our simulations. In such a dynamic system, applying standard restraints would result in trapping the structure in one (potentially unrealistic high-energy) structure or blowing up the structure in the unrestrained regions. Similarly, applying many NOEfixes would effectively freeze the structure.^53^ For this reason, we have only used few NOEfixes on selected RNA-protein distances (see Methods and **Table 1**) since NOEfix allows to sample free energy surface regions with NOE distances above upper bounds. In U1 SL3 and hnRNPA2/B1 FUS systems, three and four NOEfix-curated NOEs showed no violations in the production ensemble **S** while the remaining three were violated despite the use of NOEfix (**Table 1**).

For the sake of completeness, to the best of our knowledge the complexity of the system’s free energy surface, the force-field problems and the low amount of primary NMR data does not justify using any currently available methods of a rigorous integration of primary experimental data into the MD ensemble (see an example in Supporting Information).

### Comparison of NMR and MD data suggests multiple substates for the RNA-protein interface and RNA loops

Loop 5’-ends of both RNA hairpins (cf. ***Figure 1***f) are flexible and sample multiple conformations in simulations. The effect is more pronounced for the U1 SL3 loop where all experimentally measured NOEs cannot be satisfied simultaneously and where we clearly found different local substates with mutually conflicting NOEs. Reported cross-linking and immunoprecipitation (CLIP) studies showed that numerous RNA binding proteins cross link to U1 SL3 5’-end *in vivo*.^60^ Therefore, the different RNA substates we observed in our simulations could be recognized by different RNA binding proteins *in vivo*. For the hnRNPA2/B1 stem loop, we did not identify such clear local sub-states for the loop 5’-end and only two violated NOEs fluctuated slightly above the upper-bound values. Thus, the simulations suggest U1 SL3 loop to be more dynamic than the shorter hnRNPA2/B1 loop.

The bases at the 5’-end of the RNA loop generally sample reversible extrusions to the solvent. Bulging out is often facilitated by stacking against arginine residues. Sometimes, the bulging involves just one nucleobase while sometimes two stacked bases bulge out together. Notably, multiple states with bulge-out bases are associated with some of the rarely-reproduced FUS-U1 SL3 NOEs (see below). Generally, bulged-out bases could become accessible for other possible partners, protein or RNAs, with which bulged adenines can form, e.g., A-minor interactions.^61^ Also, bulging compacts the loop.

For the RRM-bound YNY motif, we have mainly observed base unstacking from protein in the middle position (G_16_/U_16_, see below). Coexistence of two states is supported by NMR data for hnRNPA2/B1 U_16_. For the outer (Y15 and Y17) positions, the base-protein H-bonds are generally quite fluctuating but the nucleotides retain their positions in their pockets while the NOE data do not allow us to conclude anything specific about the observed dynamics (see Supporting Information for details). Imbalanced description of H-bonding at RNA-protein interfaces is rather common in RNA-protein simulations.^62^

In the following sections, we describe RNA sub-states that could explain for the individual NOE violation at the binding interface and within the RNA loops (violation > 1 Å, **Table 1**), analyze their mutual conflicts and estimate their minimal populations when possible. Full list of NOE violations can be found in Supporting Information.

Among the violated NOE-derived distances, few clearly contradict some of the other NOEs. This indicates co-existence of different local sub-states. For such NOEs, the violations typically emerge already during the extended equilibrations (see Methods) suggesting structural strains in these regions in the starting structures. Then the simulations only rarely sample distances below the upper bound threshold. In the starting NMR structures, the distances are either satisfied or only marginally (< 0.3 Å) violated since they are restrained.

It should be noted that the ensemble of sampled RNA structures is very heterogeneous and we still could miss some important substates. Our analysis is done in “data-poor” regime and is certainly affected by the force field. Sampling of the RNA-protein interactions is even more heterogeneous and less convergent than the RNA sampling (namely the RGG tail, see below). Almost certainly, there exist plausible substates which are entirely absent in our ensembles and which could reproduce the respective signals. Further, it is possible that some of the short-living rare RNA conformations could be stabilized by RNA-protein interactions which were not sampled in our simulations, i.e., none of the simulation trajectories reached the specific combination of RNA and protein conformations.

#### Multiple positions of A_14_ in its pocket are required to explain FUS-U1 SL3 NOE signals while its simulation behavior is comparable in both complexes

We suggest that positioning and sampling of A_14_ in its shallow RRM pocket is dynamical and the measured NOE signals reflect multiple states. Several NOEs in this region were not properly reproduced in the FUS-U1 SL3 **S** ensemble (**Table 1**). ***Figure 5*** shows three positions of the A14 which help to understand the A14 dynamics. The A_14_(H3’)-Tyr325(QD/QE) and A_14_(H2)-Asn323(HB2) NOEs violated in the overall **S** ensemble are well reproduced in simulation parts with an alternative A_14_ position at the RRM surface pocket which differs from the NMR structure (***Figure 5***b). Further, we observed A_14_ sliding from the pocket as well as reversible bulging-out. Bulging-out (not shown in ***Figure 5***) was mostly driven by A_14_-Arg328 stacking (Arg328 is from RRM, loop L3). Notably, A_14_(H8)-Arg328(QD/QG) NOEs measured for FUS-U1 SL3 are clearly anti-correlated with A_14_-pocket NOEs (Supporting Information, Figure S8). These NOEs are only satisfied in trajectory parts where A_14_ is out of its pocket; both A_14_ bulged out and sliding positions would contribute to the signal. Also the A_14_(H3’)-Arg372(QG) FUS-U1 SL3 NOE was reproduced only if A_14_ departed from its pocket (***Figure 5***c). Further A_14_ dynamics in the pocket is suggested by intra-RNA A_14_(H8)-U_15_(H4’) NOE, which is violated in all simulations, except of a few very short trajectory portions. When inspecting the associated trajectory parts and NOE violations in them, only one conformation could reasonably explain the A_14_(H8)-U_15_(H4’) signal. It had A_14_ flipped in the pocket pointing H8 atom towards the protein with simultaneous A_13_ bulge-out (***Figure 5***d); A_14_ rearranged with the help of Arg328 stacking. This substate was found in two simulations only for a few dozens of ns. If MD description of this substate gives the right distance distribution, its ~40% population would be needed to satisfy the A_14_(H8)-U_15_(H4’) NOE and it would not lead to any new violations since just ~60% weight on the (remaining) **S** ensemble snapshots would be sufficient to prevent them. The substate would also reduce (but not eliminate) the A_14_(H2)-Asn43(HB2) NOE violation but increase the present A_14_(H1’)-Tyr325 NOE violations.

**Figure 5:**
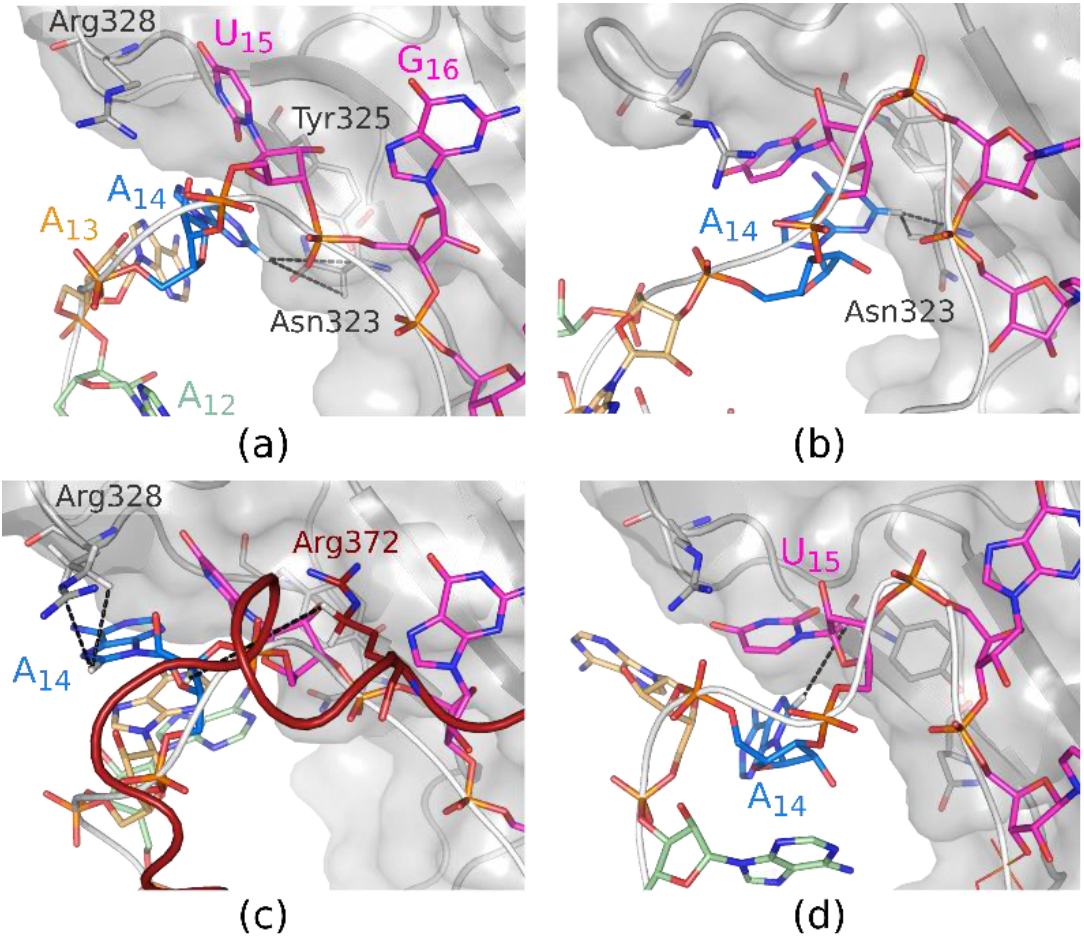
Examples of A_14_ dynamics in the FUS-U1 SL3 complex. Note that the Figure shows only part of the A_14_ dynamics and also positions of the other shown residues vary widely. (a) NMR-refined conformation closest resembling the state in (b). Note that A_14_ position in the pocket is defined by multiple RNA-RNA and RNA-protein NOEs in the NMR-refìned structures, selected NOE distances are shown by dashes. (b) Rare state producing strong NOE signal for the A_14_-Asn323 pairs. The distances in (b) are ~2 Å while NOE upper bounds are 4.5 Å. (c) A_14_ sliding in the pocket with A_13_ partially replacing it in the pocket. This and similar substates are essential to produce the Arg328 and Arg372 NOE signals to A_14_. (d) Rarely sampled substate producing the challenging A_14_(H8)-U_15_(H1’) NOE. A_14_ is flipped or rotated in the pocket while A_13_ is out. Note that restraint on this NOE may excessively push the A_14_ towards U_15_ and the protein in the NMR ensemble (a), which is identified by MD as a strained position.

Thus, to explain all the A_14_ NOEs, we suggest the presence of some conformations seen in ***Figure 5***b, some population of A_14_ out of the pocket (e.g., either sliding shown in ***Figure 5***c or bulging out) and, finally substates with flipped A_14_ position (***Figure 5***d).

The A_14_ dynamics in the FUS-hnRNPA2/B1 simulations is qualitatively similar but because the upper bound NOEs are larger, the simulations have less problems to satisfy them. The FUS-hnRNPA2/B1 system does not contain any analogous problematic intra-RNA NOE.

#### RRM-bound U_16_ of the FUS-hnRNPA2/B1 system samples two conformations in its binding site

To support the position of U_16_ in the FUS-hnRNPA2/B1 system, multiple NOEs were observed. However, U_16_-Thr286 NOEs were violated in all *main* set simulations (**Table 1**). Yet, in some trajectory parts, those NOEs are transiently satisfied (in ~1% of the **S** ensemble) at the expense of the U_16_-Phe288 NOEs. These trajectory parts show U_16_ unstacking from Phe288 and its sugar shifting relatively to the protein (**Figure 6**). The same conformation was adopted if U_16_-Thr286 NOE restraints were applied in test simulations while upon release of those restraints, U_16_ immediately slid back to its dominant conformation with Phe288 stacking. This suggests that U_16_ samples at least two positions in the binding pocket, each of the positions corresponding to different sets of NOEs. Reweighting NOEs from continuous trajectories shows that about 35% of the states could have unstacked U_16_. The NMR ensemble suggests only one U_16_ position, the stacked one, with potential local strain in the region. MD simulations might be overestimating the stacked conformation due to potential stacking over-stabilization.^6^

**Figure 6:**
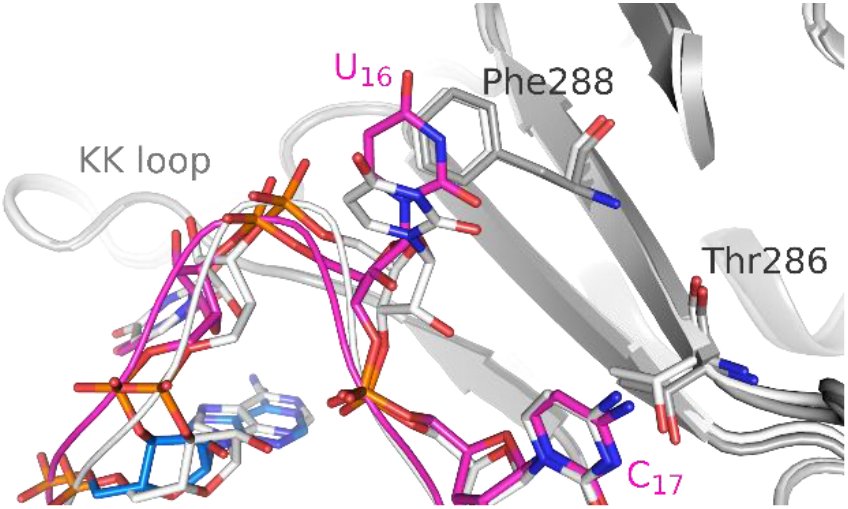
Overlay of FUS-hnRNPA2/B1 U_16_ positions. Average structure of the whole simulation **S** ensemble is shown in colors while average structure for ensemble of snapshots with at least one of the two U_16_-Thr286 NOEs satisfied is in white. RGG tail is not shown for clarity.

Also G_16_ of U1 SL3 is not rigid in this binding site and samples transient unstacking similarly to U_16_. G_16_ dynamics can be affected by the N-terminal tail of the protein, since it contacts G_16_ in the NMR structure but did not lead to intermolecular NOEs observation (see Methods). NOE data neither directly support nor rule out some population of U1 SL3 G_16_ unstacking.

#### The 5’-end of the U1 SL3 loop samples several substates

For U1 SL3 A_12_, experimentally measured NOEs support a proximity to the other nucleotides: C_11_, A_13_ and A_14_. Already during the MD equilibration, A_12_ markedly changed its position, indicating that its position was associated with a marked strain in NMR-derived structure. A_12_ samples multiple positions in the loop, most often being stacked on top of C_11_, or A_13_, or reversibly bulging into the solvent. Some of the sampled states were consistent with different NOE signals while not contributing to some other signals. Poorly reproduced in simulations were several C_11_-A_12_ NOEs and A_12_(H2)-A_14_(Q5’) contact (**Table 1**).

For the A_12_(H2)-A_14_(Q5’) contact, the simulations suggested several states that could explain this single NOE while none of them would contribute to the C_11_-A_12_ signals. The most significant ones are the following three (see also **Figure 7**):

i. State with a sharp turn in backbone between A_12_ and A_13_ with A_13_-A_14_ stacking. A_12_ is stabilized near A_14_ by stacking with RGG tail backbone. U_15_ shifts in its pocket forming the U_15_(N3-H3)-Thr326(OG) H-bond rather than the native U_15_(N3-H3)-Thr326(O) one. This state was found in three simulations with lifetimes from a few to ~200 ns.
ii. Conformation with A_12_-A_14_ stacking and A_13_ bulged out towards the minor groove. The state has a backbone conformation resembling backbone topology of RNA kink-turn motifs (the 6p-2[suite according to the standard classification^63^). This state was found to be reversibly formed in one simulation only, with maximum lifetime of 50 ns.
iii. Conformation similar to state *i* but with larger fluctuations and with A_13_ bulged out. The A_12_ position is also stabilized by RGG tail.

**Figure 7:**
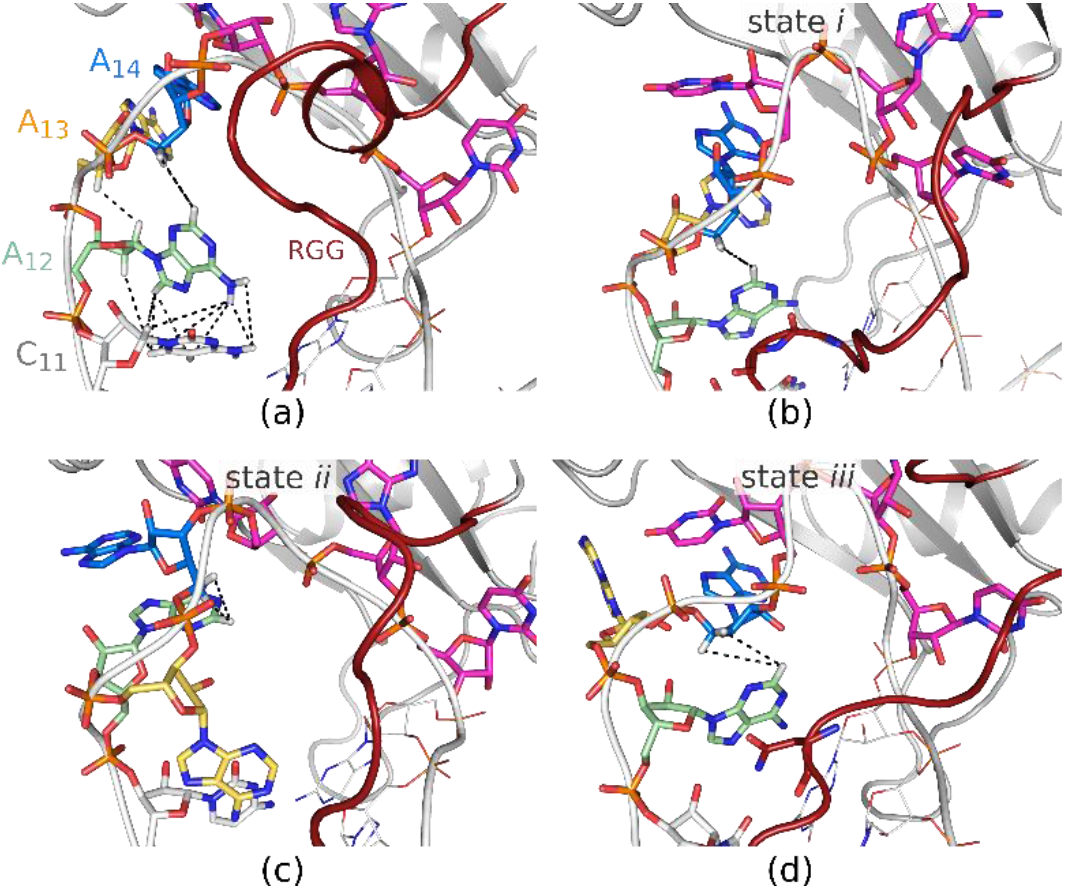
The most salient substates (*i-iii*) which could contribute to the A_12_(H2)-A_14_(Q5’) NOE; see the text for further details, (a) shows NMR-refined A_12_ position. A_12_ position is stabilized in state *i* (b) by stacking with RGG tail glycines and in state *iii* (d) by stacking with RGG asparagine; however, it was also found to be stabilized by arginine sidechains or glycines.

In order to explain the experimental data, combined population of 15-20% of continuous trajectories of the states *i* and *ii* (including fluctuations violating the distance) in the ensemble would be sufficient. In addition, some other, minor and/or unidentified structures can contribute as well. The signal could be also bolstered by some trajectory parts with progressive loss of the binding interface found in the **U** ensemble, i.e., unbound/partially unbound conformations could also contribute.

For the C_11_-A_12_ pair, five out of thirteen NOEs were significantly violated in the **S** ensemble **(Table 1Chyba! Nenalezen zdroj odkazů.**). However, it was challenging to suggest plausible states that would explain NOE signals between C_11_ and A_12_ either due to limited sampling or force-field deficiencies. There were some states reproducing the signal for one or more of the five problematic NOEs but these were either very unstable (state *j*, see below) or not capable of producing signal strong enough to fully explain the measured NOEs (states jj and jjj). The most salient substates were (see also ***Figure 8***):

*j*) State with C_11_ exposed to solvent. It is consistent with four of the violated C_11_-A_12_ NOEs and produces particularly strong signal for C_11_(H6)-A_12_(H2’) (3.3 Å below upper bound) but contradicts C_11_-G18 NOEs. This state was only found in one simulation and persisted for just ~3 ns. Subsequent simulation runs initiated from this conformation could not retain it while supporting the conformation with NOEfixes prolonged life-time of the conformation to up to 20 ns but then the structure deteriorated irreversibly.
*jj*) State with C_11_ and A_12_ unstacked and A_12_ amino group in the plane of the C_11_ base. The inclined A_12_ position was often stabilized by the RGG tail. This state satisfies the C_11_(H1’)-A_12_(H62) NOE while the A_12_ inclination contradicts the other four C_11_-A_12_ problematic NOEs. This state was found in multiple simulations with lifetimes up to 100 ns with large fluctuations.
*jjj*) State with sharp turn in RNA backbone and A_14_ shifted out of its pocket. This state satisfies the C_11_(H6)-A_12_(H2’) NOE and contradicts the other four C_11_-A_12_ problematic NOEs and A_14_-protein NOEs. It was found only in two simulations (the longer life-time was 30 ns).

**Figure 8:**
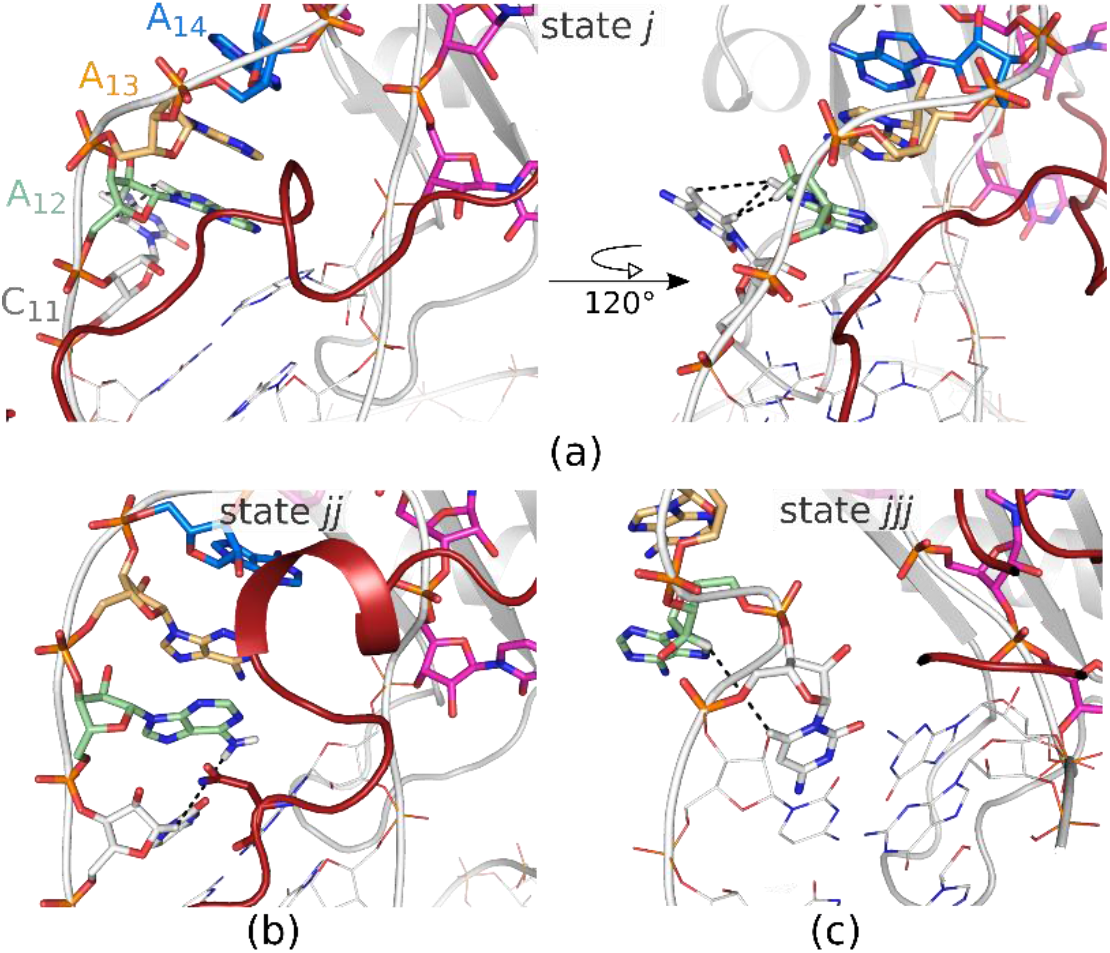
The most salient substates (*j-jjj*) which could contribute to some rare C_11_-A_12_ NOE signals; see the text for further details and **Figure 7**a for NMR-refmed position. The A_12_ position in state *jj* (b) was stabilized by stacking with RGG asparagine (shown in the Figure) or glycines, but it could also occur in absence of any obvious protein-A_12_ interaction.

States *jj* and *jjj* would have to reach populations of about 70% to eliminate the respective NOE violations in the ensemble but at the same time these states conflict other NOEs. Thus, they can contribute to the signal but cannot fully explain it solely on their own. The very rare state *j* would eliminate most of the violations if stabilized. Thus, we hypothesize that *j* and similar RNA conformations may be relevant substates. From the point of view of RNA structure *per se*, the state looks plausible. The C_11_-A_12_ nucleobases are exposed for potential stacking and H-bonding interactions and might thus be captured (stabilized) by the RGG tail. For example, RGG arginine or asparagine could H-bond to A_12_ Hoogsteen edge while being stacked with C_11_. As we emphasize throughout the text, due to enormous flexibility of the system, our sampling is quite limited and thus it is possible that the simulations simply did not find the right combination of RNA and protein conformations. Similar stabilization of solvent-exposed nucleotides by interaction with flexible protein chain was reported, for example, for Moloney murine leukemia virus RNA,^64^ though in that case, the RNA-protein binding was structurally more stable.

Finally, for intemolecular violated NOEs, one of the RNA-Met321 NOEs in both complexes gets violated in all trajectories. The simulations did not find any specific structural feature associated with some short trajectory parts (at least tens of ns long) where this NOE is reproduced.

### The RGG tail is highly disordered in simulations but remains localized in the RNA minor groove

The simulations confirm that the RGG segment is extremely flexible and remains disordered when bound to the RNA. In simulations, each RGG amino acid samples many interaction sites on RNA (***Figure 9***) resulting in dozens of different possible (transient) interactions (H-bonding, stacking and salt-bridges) with different dynamics (lifetimes up to singles of μs for arginine stacking combined with H-bonds or salt-bridge). In hnRNPA2/B1 minor groove, RGG interacts more with RNA backbone than with the bases (with roughly 2:1 ratio), while this difference is less pronounced for the stiffer U1 SL3 stem (Supporting Information, Figure S10). Some examples of RGG interactions with RNA have already been shown above (**Figure 7**a and c and ***Figure 8***b) and we include more in the Supporting Information Figure S9. With such dynamics it is not possible to do any quantitative analysis of the RGG-RNA interactions. The RGG segment is sampling mainly the RNA loop, with arginine residues often entering the loop interior and the minor-groove. In the stem groove, RGG preferentially interacts with backbone of the 3’-strand, around the 4^th^ base pairs after the loop (***Figure 9***). This sampling can be affected also by the length of the RGG tail and presence of artificial C-termini at the end of the cut tail instead of the natural continuation of the tail. Similarly to RGG arginines, Arg328 of the loop L3 also dynamically H-bonds and stacks with its nearby nucleotides.

**Figure 9:**
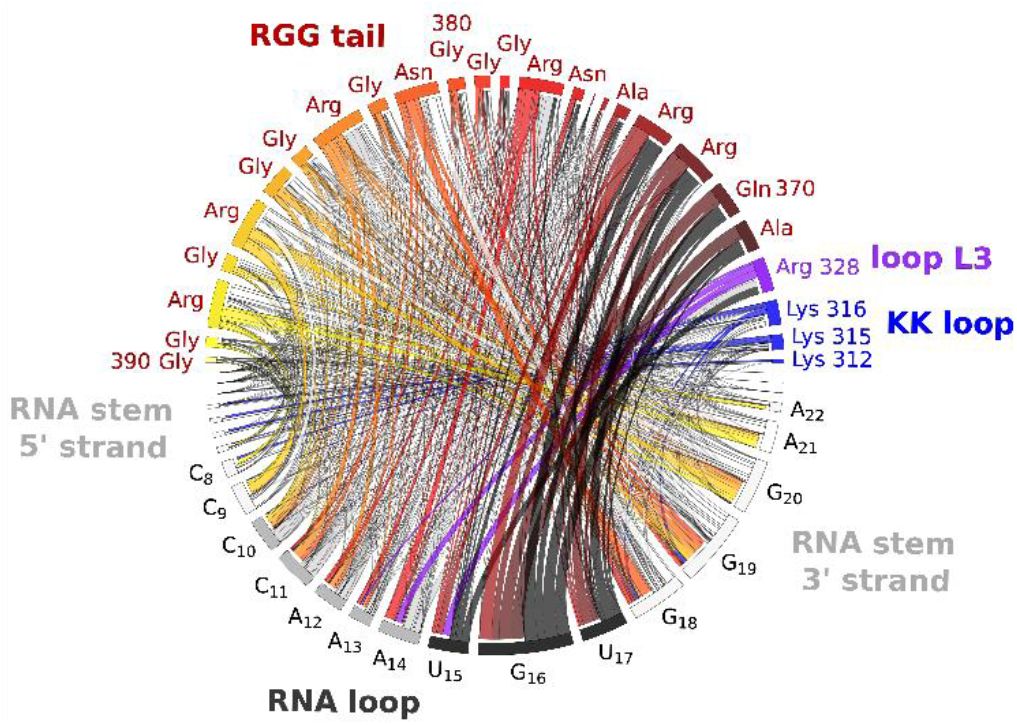
Intermolecular H-bond network map for FUS-U1 SL3 complex S ensemble. Relative widths of the lines represent number of interactions formed between the residues. Stacking interactions are also present in the system but not included in the Figure. The Figure shows that each RGG residue interacts with multiple U1 SL3 RNA residues while 3 *‘*-strand residues following the loop (G15-G20) are frequently contacted by the RGG tail. The graph was generated with Circos visualization tool. See Supporting Information Figure S11 for the FUS-hnRNPA2/B1 complex.

Experimental measurements for FUS-U1 SL3 show multiple NOE pairs between RGG glycines and RNA stem minor groove (H1’ atoms). Given the long upper-bound distance (6 Å), transient contacts between the residues are sufficient to reproduce the signals. Only for the C_8_(H1’)-Gly384(HA) pair, we did not manage to reproduce the signal in MD ensembles and the signal was lost already during the equilibration process. However, the RGG-RNA NOEs were treated as ambiguous restraints during the assignment and the initial stage of NMR structure calculation since due to the overlap of the NMR resonances of glycine and arginine residues from the RGG tail, the precise assignment of NOE signals to individual residues is challenging. The simulations show that rather Gly384, Gly386 and Gly388 are closer to C_8_ residue and would reproduce the signal. NOEs for RGG-hnRNPA2/B1 were not measured but when the U1 SL3 ones are mapped onto the respective hnRNPA2/B1 residues, most RGG-RNA signals are reproduced as well. This suggests that the RGG localization in both RNA stems could be similar.

The only stable RGG interaction which we have observed (in the *preliminary* set of simulations of the FUS-U1 SL3 complex) was Arg383 sidechain stacking on top of RNA stem with G18, accompanied by C_11_-Arg383 H-bond pairing. It caused compaction of the RGG tail (Supporting Information, Figure S17). Analogous arginine pseudo-base-pairing and stacking pattern is formed by DNA-repair proteins.^65, 66^ Minor population of this interaction cannot be ruled out based on the primary NMR data. However, it is clearly over-stabilized by the force field, as discussed in Supporting Information. In case of hnRNPA2/B1, no arginine formed an analogous permanent stacking interaction since its stem lacks the rigid GC stacking platform. More details and auxiliary simulations are provided in Supporting Information.

Localization of RGG in the minor groove in our simulations fully supports the hypothesis that RNA binding affects RGG conformational sampling and thus helps position the ZnF motif toward its exonic binding site.^26^ Yet, though the overall RGG behavior in simulations is in agreement with experimental data (disordered nature and position in the minor groove), we cannot guarantee that the ensemble and interactions details are accurate since the experimental data on RGG are scarce. Moreover, details of RGG simulation behavior are almost certainly force-field dependent.^67^ The ff14SB+SPC/E force field used in this study is known to be prone to sampling of more compact structures so the real dynamics of RGG tail could be even larger than what is observed in our simulations.

Importantly, the simulations correctly captured the principally different properties of the different RGG-RNA-binding interfaces of FUS, FMRP and SF3A1 proteins (simulations are presented in Supporting Information). While the FUS RGG remains disordered in simulations (even with the force field that supports compacted structures), the FMRP and SF3A1 RGG segments remain structured in major grooves with well-defined positions. Structuring of FMRP RGG peptide in the complex with RNA when starting from a looser conformation was reported earlier.^31^ Thus, there is in all three cases a full agreement between the simulations and experiments regarding the ordering of the RGG segment upon RNA binding. This suggests that force fields are able to correctly capture qualitative differences in behavior of different RGG-RNA binding interfaces.

### MD simulations suggest coupling among different segments and interactions across the whole FUS-RNA complex and a role for the RNA stem flexibility

In previous sections, we focused on the RNA-RRM interface, RNA loops and RGG tail. In the following paragraph, we briefly illustrate that a fully stable FUS RRM-RGG-RNA complex also requires presence of the other parts of the complex (***Figure 1***). In general, the simulation data suggest that the sequence of the stem may play a role in fine-tuning FUS-RNA molecular recognition, both by local residue-specific interactions with the RGG tail and the KK loop and by its overall position with respect to the RRM. The simulations indirectly suggest “multi-body” coupling among the different segments and interactions ranging across the whole simulated complex (cf. **Figure 1**), utilizing the natural flexibility of its different parts to optimally position all elements participating in RNA recognition. This coupling may also contribute to problems in the force-field description, since even relatively subtle inaccuracies in description of local structural elements could interfere with the overall arrangement of the complex.

The RNA stems are neutralized by the RGG tail and the lysine residues of the KK loop. MD simulations show that the stem bending due to thermal sampling allows the KK loop lysine residues to transiently contact the stem along its whole length. The positively-charged side chains of the KK loop are essential for RNA binding.^68^ When we mutated the KK loop lysines to alanines, we noticed changes in the position of the RRM with respect to the RNA hairpin and more profound losses of the YNY motif binding compared to simulations with unmodified KK loop. The loop seems to be important not only for the neutralization of the RNA stem but also for overall protein positioning with respect to the RNA hairpin. This is further supported by simulations performed using a short A_14_-G18 single strand construct derived from U1 SL3, where the experimentally determined position of U_17_ could not be reproduced unless we appended a segment of canonical A-RNA helix following U_17_ (See Supporting Information, Figure S16).

The hnRNPA2/B1 stem segment downstream the loop is dynamic and adaptable, which enables the protein (KK loop and RGG tail) to contact the stem more extensively and variably than in the case of U1 SL3 (Supporting Information Figure S12). Similar stem dynamics was also observed in auxiliary simulations of the free hnRNPA2/B1 RNA, albeit with shorter lifetimes. In contrast, the only flexible element in the U1 SL3 stem is the C24 bulge. To see how the FUS binding may cooperate with flexibility of the U1 SL3 stem, we have performed simulations of the isolated stem with and without the bulge. By comparing both simulation, it seems that the C24 bulge increases the flexibility and allows for sampling conformations closer to the FUS RRM, mainly to the KK loop (Supporting Information, Figure S15). Even transient interactions between the stem and KK loop might contribute to the binding albeit it is fair to admit that C24 is already far from the main RNA-protein interface. Flexibility of the RNA stem thus may have some role in protein binding, in lines with several recent reports suggesting a role for sequence-dependent shape and flexibility of A-RNA (or hybrid) duplexes in several molecular recognition processes^69–71^ and for single-nucleotide bulges in increasing flexibility and affinity for protein to A-RNA.^72^ For more details and auxiliary simulations see the Supporting Information.

## Concluding remarks

Biomolecular complexes often comprise segments with different degrees of flexibility. Studies of flexible molecules, however, pose a great challenge for structural experimental methods. Thus, it is tempting to combine or complement them by MD simulations, which aim to describe dynamical molecular ensembles. However, flexible parts of molecular complexes are also very challenging for modelling methods, due to force-field and sampling limitations, as well as paucity and ambiguity of the primary experimental data. The conformational space of RNA-protein complexes is nowadays commonly studied by MD simulations. However, there is no consensus on how reliable such studies are. Many studies primarily present simplified analyses based on short standard or enhanced-sampling simulations, lacking a rigorous assessment of the agreement with the experimental structures. Without reporting structural details and convergence analyses, the results are difficult to fully comprehend. It was shown that while some RNA-protein complexes can be simulated stably, other systems deteriorate progressively along the simulation trajectories.^5, 6, 31^

We used extended atomistic explicit-solvent MD simulations to complement NMR spectroscopy experiments recorded on two FUS RNA-protein systems. The FUS-RNA complexes contain well-structured RRM domain with low sequence specificity for the RNA binding, disordered RGG tail, an RNA hairpin with 6 or 7 nucleotide loops and an RNA stem (**Figure** 1). Our study provides several suggestions which extend the information from the experimental data but also demonstrate methodological complexity of MD studies of such complex RNA-protein interfaces.

Most RNA-RRM structures described to date possess structurally well-defined interfaces with short ssRNAs and are quite well described by MD simulations. However, the FUS-RNA complexes turned out to be challenging. Despite our efforts to stabilize the binding via several system-specific force-field adjustments, we have often observed progressive distortions of the RNA-protein interface inconsistent with experimental data (***Figure 3***). The simulated ensemble is also clearly unconverged. Different parts of the system sample multiple conformations on their own which creates an overwhelming spectrum of combined substates of the complex as a whole. The individual simulations sample non-equivalent ensembles, so that the trajectory set is very heterogeneous (Supporting Information, Table S4 and Figure S7). On the other hand, the simulations clearly indicate conflicts in the starting experimental structures, suggesting that these molecular complexes cannot be fully represented by a single static structure (***Figure 4*** and “Comparison of NMR and MD data suggests multiple substates for the RNA-protein interface and RNA loops” section). It justifies the use of simulations to unravel the dynamics.

Since the FUS-RNA systems behave very differently from RRM-RNA systems we studied previously, we had to adopt a completely different strategy to perform and analyze the simulations (***Figure 2***). First, we applied a set of mild system-specific force-field modifications to slow down severe distortions. Second, we split the overall simulation ensemble into two parts. The **S** ensemble includes parts of the trajectories which preserve significant similarity with the starting structures and reasonable agreement with the primary experimental data. We analyzed substates in the **S** ensemble upon assumption that the trajectories, although being affected by simulations problems, do reflect also real properties of the system. We identified sampled structural features whose combination could help to reconcile the NOE data. The remaining part of the simulations (the **U** ensemble) was discarded. The analyses of the **S** ensemble lead to the following suggestions. The FUS systems exist as dynamical ensembles. The hnRNPA2/B1 U_16_ residue samples two states (stacked and unstacked) on the RRM (**Figure 6**). U1 SL3 RNA loop is more dynamic than hnRNPA2/B1 and residues C_11_-A_14_ sample multiple local substates, including bulging-out (**Figure 5**, **Figure 7** and **Figure 8**). The flexible adenines are well poised to interact with other molecules. The RGG tail is localized in the minor groove but remains disordered, sampling astonishing range of transient interactions with RNA, including interactions stabilizing rare RNA substates. The simulations indicate presence of some long-range couplings among the different elements contributing to the RNA-protein recognition which can lead to some allosteric communication throughout the system. For example, the RNA stem-KK loop interactions affect the position of the RRM with respect to the RNA and the sequence-dependent flexibility of the RNA stem could fine-tune the binding.

In summary, our work provides insights into the potential richness of the structural ensemble of the FUS-RNA systems which is not fully apparent from the experimental data but also highlights methodological challenges in studies of RNA-protein complexes.

## Supporting information

Starting Structures and NOE violations

## Acknowledgements

This work was supported by the Czech Science Foundation project number 20-16554S (MK and JS) and the project SYMBIT reg. number: CZ.02.1.01/0.0/0.0/15_003/0000477 financed by the ERDF (PP and JS). SC is supported by INSERM and by La Ligue contre le Cancer though the grant APPRNA 2021.LCC/SeC. The authors would like to thank Prof. F.H.T. Allain for helpful discussions.

## Supporting Information

Starting structures and sampled states (ZIP)

Table of NOE violations (xlsx)

List of simulations, HBfix/NOEfix biasing potential, setup for FMRP and SF3A1 RNA-RGG simulations, selection of the optimal force field, list of **S** to **U** ensemble transitions, example of a disrupted trajectory, ensemble reweighting, comparison of structures reached at ends of individual simulations, NOE violations, cross-correlation matrix for NOEs, U_15_-protein H-bond network instabilities in MD, further analyses of RNA-RGG interactions, analyses of stem flexibility and KK loop binding, excessive arginine stacking in FUS-U1 SL3 complex (PDF)

## Notes

### Competing Interest Statement

The authors have declared no competing interest.

