## Supplementary Information for "Conformational Heterogeneity of RNA Stem-Loop Hairpins Bound to FUS RNA Recognition Motif with Disordered RGG Tail Revealed by Unbiased Molecular Dynamics Simulations"

by

Pavlína Pokorná,[1,2] Miroslav Krepl,[1] Sébastien Campagne[3] and Jiří Šponer[1,*]

[1] Institute of Biophysics of the Czech Academy of Sciences, Královopolská 135, 612 65 Brno, Czech Republic; [2] National Centre for Biomolecular Research, Faculty of Science, Masaryk University, Kamenice 5, 625 00 Brno, Czech Republic; [3] Inserm U1212, CNRS UMR5320, ARNA Laboratory, University of Bordeaux, 146 rue Léo Saignat, 33076 Bordeaux Cedex, France

### SI contents

### List of simulations

Table S 1: List of simulations.

| system | starting structure[a] | total sampling [μs][b] | sampling used in the **S** ensemble [μs][c] | force field and modifications |
|---|---|---|---|---|
| | | ***main* simulation set** | | |
| FUS-U1 SL3 | Equilibrated NMR structure 6SNJ, model 1 | 3x10+13x5+8x3 | 5x5, 1x2.5 | as specified in the Main text, Methods |
| | MD snapshot from stable trajectory part with well-reproduced YNY-RRM binding and $A_{12}$-$A_{13}$-$A_{14}$ triplestack | 2x5 | 1x3 | |
| | MD snapshot from stable trajectory part with well-reproduced YNY-RRM binding and $A_{13}$ bulge out | 2x5 | 1x2 | |
| | MD snapshot from another stable trajectory part with well-reproduced YNY-RRM binding and $A_{12}$-$A_{13}$-$A_{14}$ triplestack | 2x5 | - | |
| FUS-hnRNPA2/B1 | Equilibrated NMR structure 6GBM, model 1, snapshot at 100 ns from the RGG equilibration run – see Methods | 2x10+5x5 | 2x10+1x5 | |
| | Equilibrated NMR structure, snapshot at 200 ns from the RGG equilibration run | 1x10+6x5 | 1x10+3x5+2x2.5 | |
| | Equilibrated NMR structure, snapshot at 300 ns from the RGG equilibration run | 5x5 | 2x5 | |
| | Equilibrated NMR structure, snapshot at 400 ns from the RGG equilibration run | 5x5 | 4x5 | |
| | | ***preliminary* simulation set** – listed are only simulations added to U1 SL3 **S** ensemble | | |
| FUS-U1 SL3 | Equilibrated NMR structure, model 1 | 4x5 | 1x3 | OL3+ff19SB+OPC, no HBfixes and NOEfixes used, Arg383-water interactions not modified |
| | | 4x5 | 2x3 | OL3+ff14SB+SPC/E+cufix, no HBfixes and NOEfixes used, Arg383-water interactions not modified |

**auxilliary simlations**

| system | starting structure | total sampling [μs][b] | force field and modifications |
|---|---|---|---|
| U1 SL3 free RNA stem | cut out from the NMR structure | 1x0.5 | as specified in the Main text, Methods |
| U1 SL3 free RNA stem without C24 bulge | build with Nucleic acid builder of Amber | 1x0.5 | |
| FUS-U1 SL3 complex without C24 bulge | Equilibrated NMR structure combined with Nucleic acid builder model of the stem | 1x1+2x0.5 | |
| hnRNPA2/B1 free RNA | cut out from the NMR structure | 3x3+1x2.5+1x1 | |
| $C_{11}$-$A_{12}$ NOEfixes | MD snapshot with the rare $C_{11}$-$A_{12}$ NOE satisfied - state *j* in Main text Figure 8 | 2x1 | Additional NOEfixes on $C_{11}$(H1′)-$A_{12}$(H8), $C_{11}$(H6)-$A_{12}$(H2′), $C_{11}$(H5)-$A_{12}$(H8) |
| | | 2x0.5 | Additional NOEfixe on $C_{11}$(H6)-$A_{12}$(H2′) |

| | | | |
|---|---|---|---|
| FUS-U1 SL3, KK loop lysines to alanines | Equilibrated NMR structure, lysines mutated with parmed | 10x5 | |
| FUS-hnRNPA2/B1, KK loop lysines to alanines | Equilibrated NMR structure, lysines mutated with parmed | 4x5 | |
| FUS-hnRNPA2/B1 | MD snapshot with $U_{16}$-Phe288 unstacked | 2x3 | additional $U_{16}$-Thr286 NOE restaints |
| | Last snapshot of one of the simulations from the row above | 1x0.5 | restraints released |
| Stem swap, system A[d] | | 7x5 | |
| Stem swap, system B[d] | | 7x5 | |
| Stem swap, system C[d] | | 7x5 | |
| Stem swap, system D[d] | | 7x5 | |
| FUS-U1 SL3 complex with ssRNA | Equilibrated NMR structure with keeping only nucletides $A_{14}$-$G_{18}$ and RRM | 4x1 | RNA-protein vdW interaction weakened by scaling ε of selected atom pairs by 0.75 |
| FUS-U1 SL3 complex with ssRNA+helix construct | Equilibrated NMR structure with keeping only nucletides $C_8$-$C_{10}$+$A_{14}$-$G_{20}$ and RRM | 4x1 | RNA-protein vdW interaction weakened by scaling ε of selected atom pairs by 0.75 using the stafix rescaling scheme[1] |
| FMRP | X-ray structure 5DEA | 3x5 | OL3+ff14SB+SPC/E |
| FMRP | | 3x5 | OL3+ff14SB+SPC/E+cufix |
| FMRP | | 4x5 | OL3+ff99SB+OPC |
| SF3A1 | X-ray structure 7P0V with RRM and RNA loop removed | 2x5 | OL3+ff14SB+SPC/E+cufix |

[a] coordinates of the starting structures are included in Supporting Information zip file.
[b] number of simulations multiplied by the length of each simulation, i.e. 3x5 means three simulations each 5 μs.
[c] initital 100 ns were always discarded from the analyses. The decision whether the simulation is still in the **S** ensemble or already in the **U** ensemble was made each 0.5 μs based on a careful analysis of a number of factors (see Methods).
[d] see *Figure S 14* for specification of the systems.

### HBfix/NOEfix biasing potential

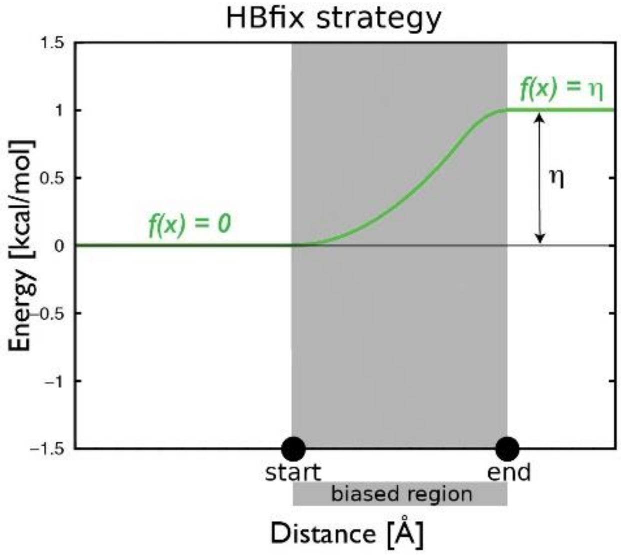

Figure S 1: Schematic representation of HBfix (NOEfix) strategy. The biasing potential (green line) is active only within the biased region and is constant outside of it. See ref. [2] for details.

### Setup for FMRP and SF3A1 RNA-RGG simulations

For simulations of RGG motif of FRMP (residues 527-544) bound to sc1 RNA we used the PDB ID 5DEA[3] structure (X-ray, 2.8 Å resolution). For simulations of SF3A1 RGG (residues 787-791) bound to stem of U1 SL4 we used PDB ID 7P0V structure (X-ray, 2.8 Å resolution)[4] from which we removed the RRM domain and RNA tetraloop. The U1 SL4 RNA helix was capped by GCG pairs at both ends. Used force fields are specified in *Table S 1*; the same equilibration and simulation protocols as described in the Main text (except for the extended NMR-restraint equilibration) were used.

### Selection of the optimal force field

In the *preliminary* simulation set we have observed spurious long-living interactions of the RGG tail arginine residues with anionic amino acids and RNA, which were notably reduced when we applied the cufix correction using published parameters.[5, 6] We have also considered OPC water model (instead of SPC/E) along with the ff19SB protein force field, since the OPC water is also supposed to prevent excessive solute self-interactions by enhancing solute-water interactions. Both OL3+ff14SB+cufix+SPC/E and OL3+ff19SB/OPC force-field versions resulted in reduction of excessive protein-stem interactions with respect to OL3+ff14SB+SPC/E to a similar extent (*Figure S 2*). Since the core of the studied RNA-protein interface consists of RRM-RNA recognition, we did not consider any force-fields developed specifically for intrinsically disordered proteins. We made a short test combining OL3+ff19SB+OPC with the cufix which, however, resulted in a loss of the binding (data now shown; note that the cufix parameters were not optimized for the OPC water model).

To assess possible side-effects that cufix could have on RNA-protein interfaces, we have also run simulations of the FMRP-RNA complex with structured RGG segment. FRMP (Fragile X mental retardation protein)-RNA complex has a well-defined RNA-protein interface, which has been characterized by both NMR[7] and X-ray crystallography[3] (*Figure S 3*A). The X-ray structure revealed more contacts between RNA and protein than originally suggested by NMR. We have performed FMRP-RNA simulations in order to test the effect of different force-field variants on structurally well-defined RNA-RGG interface. Our FMRP-RNA simulations (at least 3x5 μs per each force field combination, see *Table S 1*) were initiated from the X-ray structure (PDB ID 5DEA, see above).

Only minor differences among the used force-field variants were observed. The OL3+ff19SB+OPC force field showed tendencies to unbind the N-termini, which is documented by loses of the $U_{28}$(O2′)-Asn8(N-H) H-bond (*Figure S 4*). Simulations with OL3+ff14SB+SPCE with or without cufix correction show very comparable results regarding the interface H-bonds. OL3+ff14SB+SPCE and OL3+ff19SB+OPC simulations moreover resulted in some U3-Arg8 NOE violations while OL3+ff14SB+SPCE+cufix variant did not (*Table S 2*). Overall, we concluded that the cufix correction decreases excessive non-specific RNA-protein binding in the FUS-RNA system without corrupting well-defined native contacts in the FMRP-RNA complex. Based on these test runs, simulations in the *main* set have been carried out with the OL3+ff14SB+cufix+SPC/E force-field combination.

We however note that our benchmark is not exhaustive and the results can be affected by the limited sampling and the fact that we have initiated the simulations from already bound native state.

All FMRP simulations with all force fields show Arg10 shifting its sidechain from the crystal structure position that leads to change in the local H-bond network (*Figure S 3*B and C). The MD Arg10 conformation is neither supported not ruled out by detected NOE signals. Further, two H-bonds formed by RGG peptide N-terminal Arg8 are lost in all simulations (*Figure S 4*). However, the respective part of the RGG peptide is expected to be more dynamic and its structure in X-ray experiment is clearly affected by direct contacts with another crystal unit. Regarding the NOE distance violations, all simulations show high agreement with the NMR data. Percentages of satisfied NOEs in individual trajectories range from 98 to 97%, which is even slightly better than agreement of the starting X-ray structure with the NOE (95%, note that different experimental conditions can lead to different structures in X-ray and NMR). Most of the NOEs violated in simulations are the same ones as violated in the X-ray structure. In summary, the simulations show very good performance for the FRMP system; the differences between the X-ray and NMR are discussed also in the experimental study.[3]

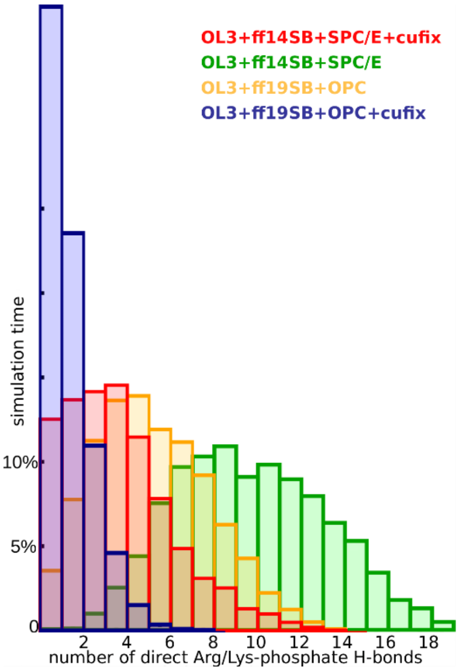

Figure S 2: RNA-protein salt-bridge propensity in FUS-U1 SL3 simulations with different force fields. Data from 2x5µs preliminary set simulations were used to produce the graph.

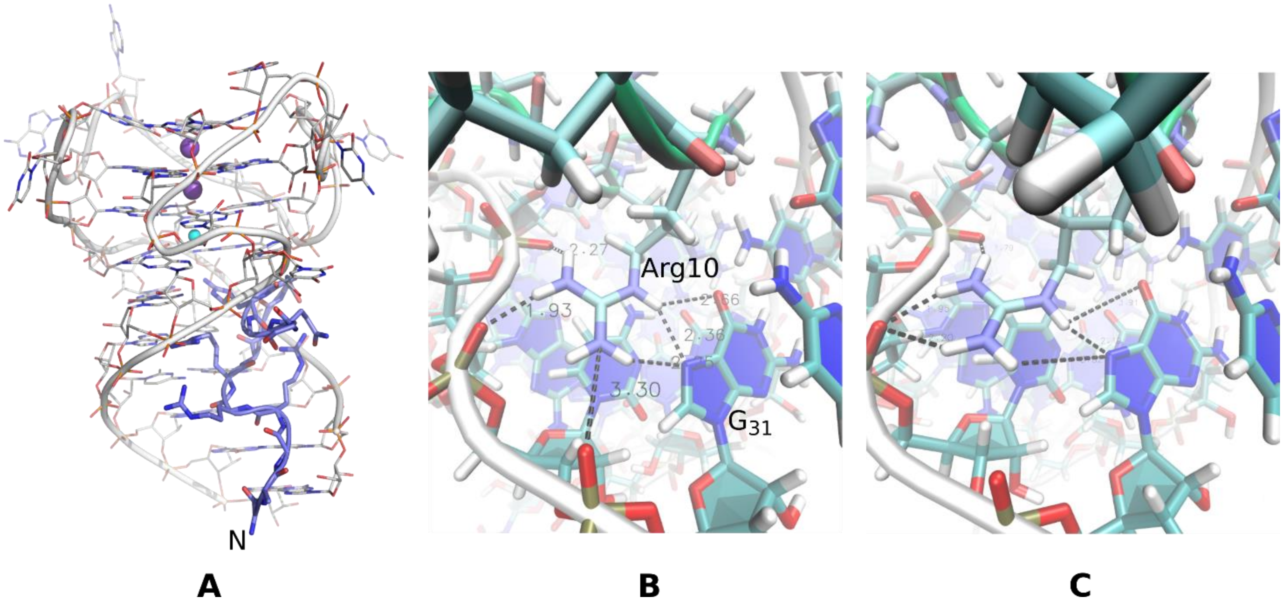

Figure S 3: **A** Structure of the FMRP peptide bound to RNA, PDB ID 5DEA. **B** Interaction network of $Arg_{10}$-G31 seen in X-ray structure and **C** in simulations.

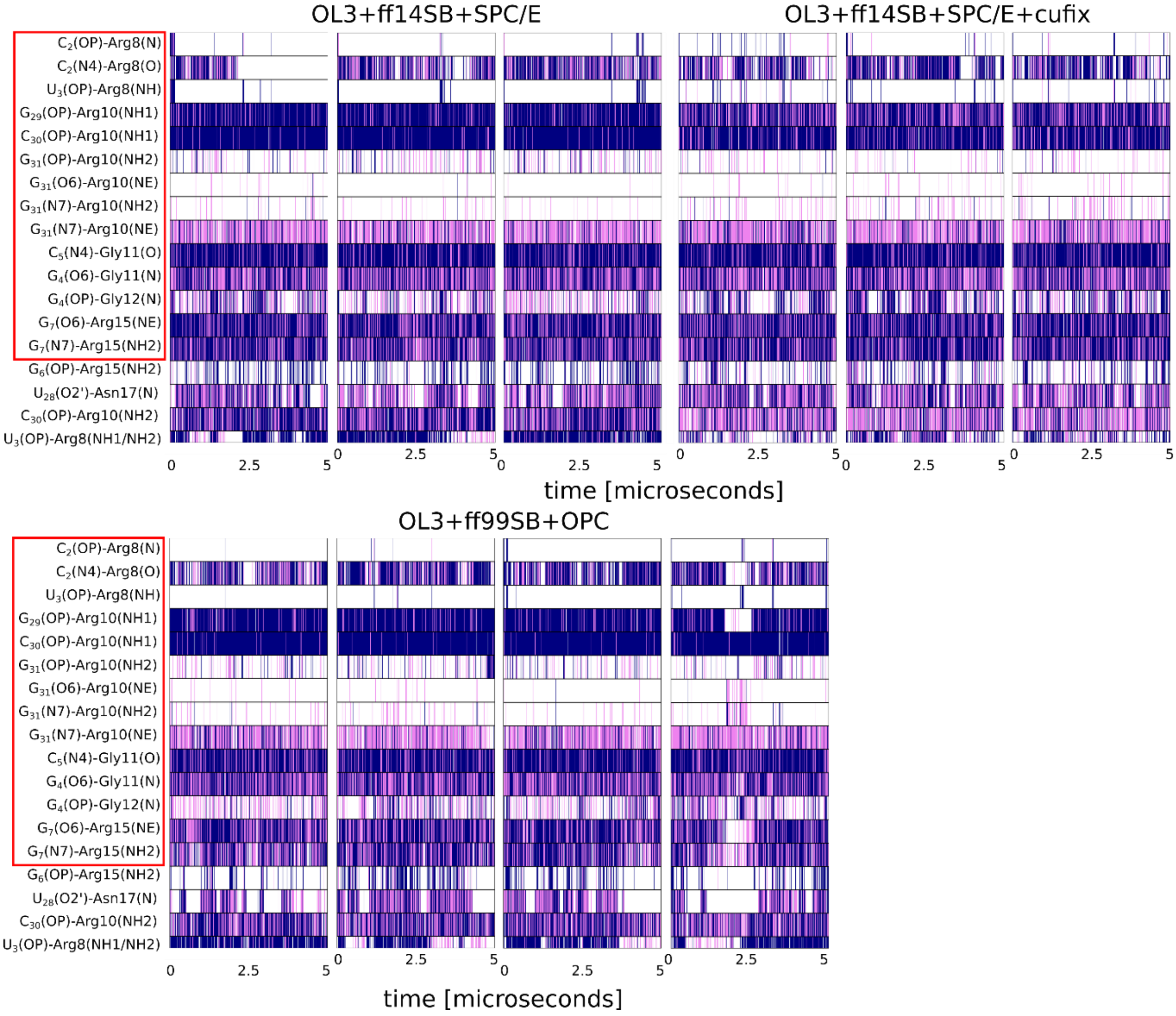

Figure S 4: RNA-protein H-bond developments in FMRP simulations with different force fields. H-bonds in red rectangles are present in X-ray structure PDB ID 5DEA. Color coding in the histograms is: blue = strong H-bond, violet = weak H-bond, white = no H-bond. The distinction is done based on following geometry criteria: strong H-bond = acceptor-donor distance < 3.0 Å and angle > 135°, weak H-bond = acceptor-donor distance < 3.5 Å and angle > 125°.

Table S 2: NOE violations in RNA-FMRP simulations.

| | number of NOEs violated in three 5-μs trajectories | | | |
|---|---|---|---|---|
| force field | RNA-protein | protein-protein | RNA-RNA | total |
| OL3+ff14SB+SPCE | 6, 6, 7 | 1, 1, 1 | 3, 3, 3 | 10, 10, 11 |
| OL3+ff14SB+SPCE+cufix | 5, 5, 4 | 1, 1, 1 | 3, 3, 3 | 9, 9, 8 |
| OL3+ff19SB+OPC | 8, 7, 7 | 1, 1, 1 | 3, 3, 3 | 12, 11, 11 |

### List of S to U ensemble transitions

Table S 3: Disruption events in 3-μs simulations from the *main* set of simulations; the simulations in the Table are numbered (ordered) according to the time of the disruption.

| disrupted simulation number | ~ time of disruption event start (ns) | description of the initial disruption event |
|---|---|---|
| | FUS-U1 SL3 complex | |
| 1 | 45 | $U_{15}$ unbinding, followed by $U_{17}$ unbinding |
| 2 | 60 | $U_{15}$ unbinding (via stacking with Tyr325) |
| 3 | 70 | irreversible changes in RNA loop |
| 4 | 110 | KK loop binding to different part of the stem and binding of the stem end to RRM, $U_{15}$ unbinding (via stacking with Tyr325), followed by U17 unbinding |
| 5 | 150 | irreversible changes in the RNA loop |
| 6 | 400 | $U_{15}$ unbinding irreversible |

| | | |
|---|---|---|
| 7 | 450 | $U_{15}$ lost followed by KK loop unbinding and irreversible changes in the RNA loop |
| 8 | 590 | Disruption of the interface around $A_{14}$, irreversible changes in the RNA loop |
| 9 | 600 | $U_{15}$ unbinding followed by $G_{16}$ unbinding |
| 10 | 800 | strong RNA stem end binding to the protein |
| 11 | 820 | RGG stably bound to RRM with no sign of reversibility |
| 12 | 1200 | $G_{16}$ unbinding |
| 13 | 1400 | $U_{15}$ unbinding followed by loss of $C_{17}$ and $G_{16}$ binding |
| 14 | 1800 | strong RNA stem end binding to the protein |
| 15 | 2100 | disruption of the interface around $U_{15}$, irreversible changes in the RNA loop |
| 16 | 2200 | $G_{16}$ and $U_{17}$ unbinding |
| 17 | 2400 | $U_{15}$ unbinding, irreversible changes in the RNA loop |
| | | FUS-hnRNPA2/B1 complex |
| 1 | 50 | $U_{16}$ unbinding |
| 2 | 60 | strong RNA stem end binding to the protein |
| 3 | 90 | irreversible changes in the RNA stem |
| 4 | 700 | irreversible changes in the RNA stem followed by irreversible changes in the RNA loop |
| 5 | 1000 | $U_{16}$ unbinding |
| 6 | 1100 | strong RNA stem end binding to protein |
| 7 | 1600 | $U_{15}$ unbinding followed by irreversible changes in the RNA stem |
| 8 | 2700 | irreversible changes in the RNA stem followed by $U_{16}$ unbinding |
| 9 | 2900 | $U_{16}$ unbinding followed by $U_{15}$ unbinding |
| 10 | 3000 | $U_{16}$ unbinding followed by $C_{17}$ unbinding |

### Example of a disrupted trajectory

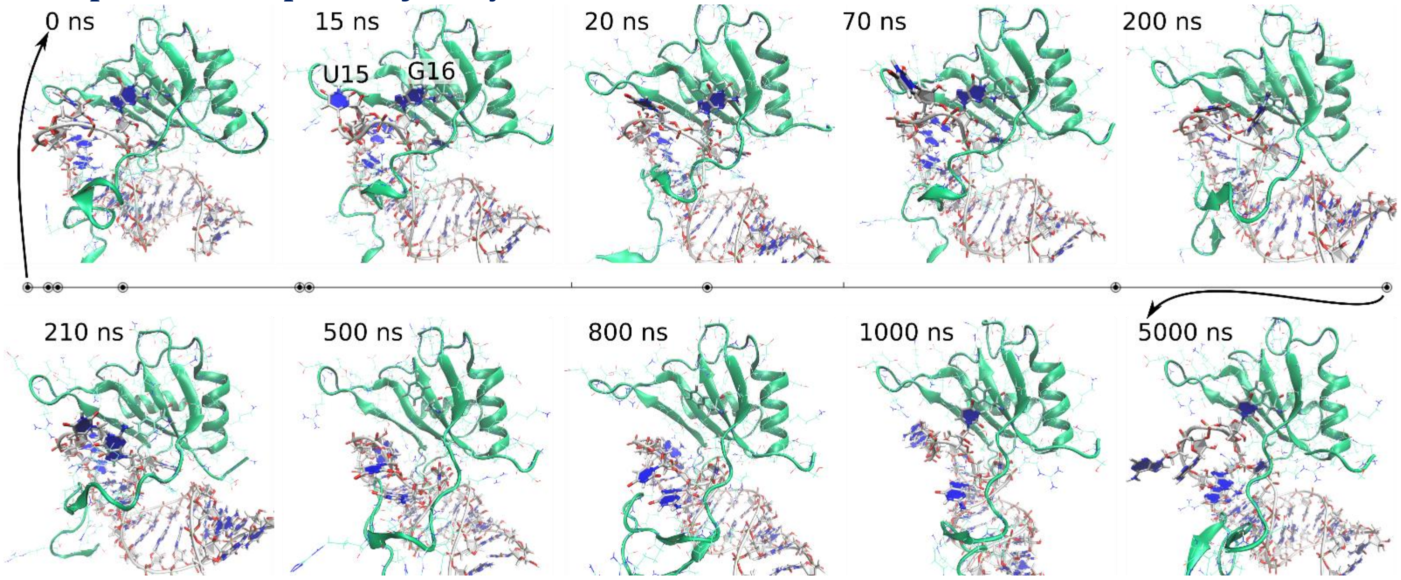

Figure S 5: Example of disruption of FUS-U1 SL3 RNA-RRM interface in one simulation from the *preliminary* simulation set. Timeline of the snapshots is shown by points on the line in the middle of the Figure. At 70 ns we see unbinding of $U_{15}$, followed by $G_{16}$ unbinding at ~200 ns and subsequent loss of the interface. At ~1000 ns $U_{15}$ binds to the $G_{16}$ pocket and the simulation retains this state till 5 μs.

### Ensemble reweighting

The Maximum entropy (ME) reweighting was done for the U1 SL3 **S** ensemble. We have used the ME code by S. Bottaro available on GitHub (https://github.com/KULL-Centre/BME) using θ=20 and all intra-RNA and RNA-protein NOEs (153 pairs). The automated reweighting resulted in $\chi^2$ drop from 0.42 to 0.18 while $\chi^2$ and number of violations after reweighting were not affected even when also the protein-protein NOEs were added to the set. The relative Kish size calculated as:

$$K_{eff} = \frac{1}{N}\frac{(\sum w_i)^2}{\sum (w_i)^2}$$

is 0.20, suggesting that only 20% of original frames significantly contribute to the reweighted ensemble. Thus the original and reweighted ensembles do not overlap well. Importantly, even though the total number of violations was lowered in our ensemble after ME reweighting, for some NOEs the lowering was only due to a small number of snapshots or even a single snapshot (*Figure S 6* A and B). Including also **U** ensemble trajectory portions to the analysis further reduces (as expected) $K_{eff}$ to 0.07. In comparison, in another study[8] where reweighting was successfully applied to RNA hairpin the $K_{eff}$ was ~0.70. Thus, the automated reweighting does not resolve the conflicts convincingly and its outcome is actually similar to our results obtained by the manual monitoring of the trajectories. Several of the NOEs can be satisfied only in very rare trajectory portions, as described in the Main text. *Figure S 6* shows an example of a search for rare sub-states explaining particular violated NOEs using our manual search for suitable sub-states as well as using ME. It is evident that both approaches find the same substates.

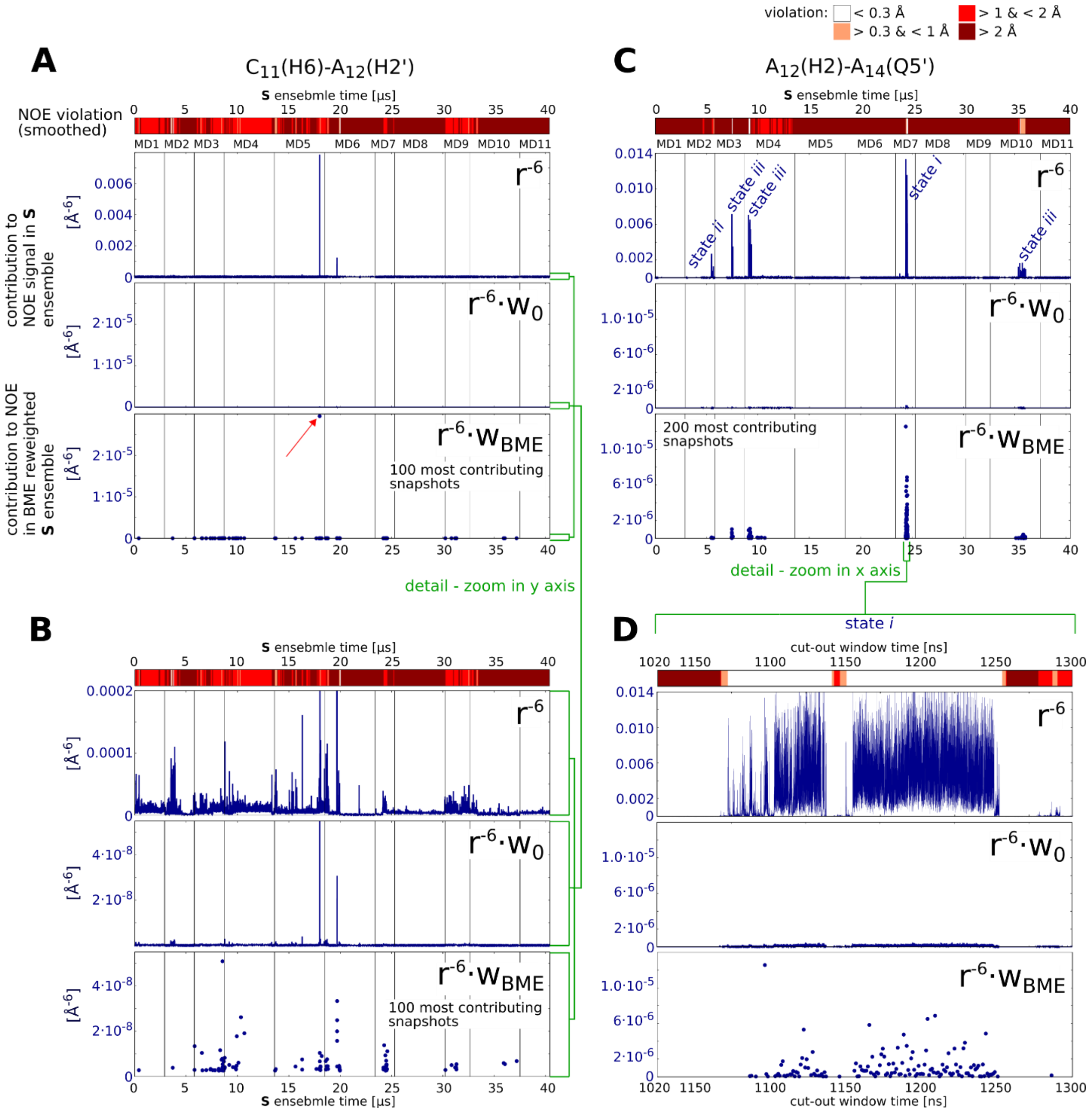

Figure S 6: Example of NOE violation analysis and identification of relevant sub-states for two heavily violated NOEs in FUS-U1 SL3. The uppermost reddish horizontal color bars in Figures **A-D** show NOE violations along the trajectories with 10 ns window smoothing. The top blue graphs show contribution of individual snapshots to the overall NOE signal, the middle show the same data multiplied by weight $w_0$ ($w_0$= 1/N = 0.000025; N=40 400 is the total number of snapshots in the **S** ensemble), while the bottom use the $w_{BME}$ weights after the BME reweighting. **A** the $C_{11}$(H6)-$A_{12}$(H2′) interaction (upper bound 4.8 Å, the closest snapshot 2.3 Å, $w_{BME}$ of the closest snapshot 0.004) monitored over the whole **S** ensemble; **B** the same with finer scales (see the green lines on the right part of the graphs) on the y-axes for better visualization. **C** the $A_{12}$(H2)-$A_{14}$(Q5′) interaction (upper bound 3.2 Å, the closest-distance snapshot 2.04 Å, $w_{BME}$ of the closest-distance snapshot 0.0009) monitored over the whole **S** ensemble. States *i-iii* are described in the Main text (Figure 7). **D** is the same as **C** but shown for the 1020 to 1300 ns period of the MD7 simulation. Graphs (**A-C**) are shown with 1 ns step while graph **D** uses 10 ps discretization, using thus N=4 040 000 in the calculation. For the $C_{11}$(H6)-$A_{12}$(H2′) NOE the BME-reweighted ensemble is heavily dominated by a single snapshot with very strong signal (marked with red arrow) while for the $A_{12}$(H2)-$A_{14}$(Q5′) there are more snapshots selected by BME that significantly contribute to the signal (see the Main text for more details). The gray vertical lines separate the individual trajectories.

### Comparison of structures reached at ends of individual simulations

Table S 4: εRMSD[9] values for U1 SL3 RNA loop (C10-G18) average structures from the last 50 ns of eleven individual simulations from the FUS-U1 SL3 **S** ensemble. Highest and lowest εRMSD values are in bolt. Note that simulation 11 was initiated from a snapshot from simulation 8. εRMSD was calculated using plumed.[10] The data show that the final structures of the RNA loop are very diverse; εRMSD is considerably more sensitive to structural differences than coordinate RMSD.

| MD # | 2 | 3 | 4 | 5 | 6 | 7 | 8 | 9 | 10 | 11 |
|---|---|---|---|---|---|---|---|---|---|---|
| 1 | 1.21 | 0.78 | 1.08 | 0.70 | 0.88 | 0.88 | 0.71 | 1.01 | 0.80 | 0.67 |
| 2 | | 1.12 | 1.09 | 1.27 | 1.11 | 0.98 | 1.23 | 1.14 | 1.02 | 1.24 |
| 3 | | | 0.69 | 0.86 | 1.03 | 0.63 | 0.75 | 1.15 | 0.96 | 0.76 |
| 4 | | | | 1.11 | 1.05 | 0.77 | 0.97 | 1.21 | 1.09 | 0.98 |
| 5 | | | | | 0.81 | 0.99 | 0.56 | 1.26 | 0.84 | **0.53** |
| 6 | | | | | | 1.09 | 0.76 | **1.33** | 0.88 | 0.77 |
| 7 | | | | | | | 0.86 | 1.02 | 0.78 | 0.85 |
| 8 | | | | | | | | 1.18 | 0.78 | 0.18 |
| 9 | | | | | | | | | 1.18 | 1.17 |
| 10 | | | | | | | | | | 0.76 |

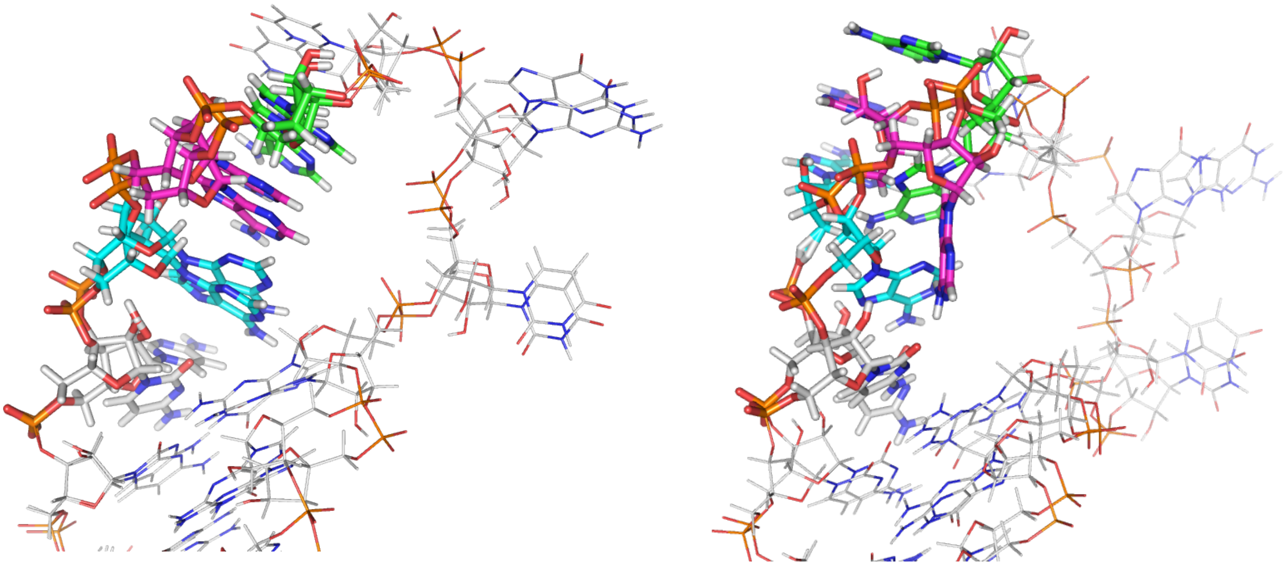

**A** **B**

Figure S 7: Comparison of the two structures from *Table S 4* with lowest (**A**) and highest (**B**) εRMSD values. $C_{10}$-$A_{14}$ nucleotides are shown as sticks and colored in a following way: $C_{10}$ white, $A_{11}$ cyan, $A_{12}$ magenta and $A_{14}$ green. The structures were aligned with respect to the whole RNA.

### NOE violations

Table S 5: NOE violations in the **S** ensembles. Full list is provided in NOEviols.xlsx file.

| type of NOE | % of NOE violated > 0.3 Å in the **S** ensemble | % of NOE violated > 1.0 Å in the **S** ensemble |
|---|---|---|
| | FUS-U1 SL3 complex | |
| RNA-protein | 13 | 4.7 |
| RNA-RNA | 6.2[a] | 2 |
| protein-protein | 16 | 6.4 |
| | FUS-hnRNPA2/B1 complex | |
| RNA-protein | 21 | 11 |
| RNA-RNA | 26[b] | 3 |
| protein-protein | 11 | 3 |

[a] mostly in the loop region.
[b] mostly in the stem region.

### Cross-correlation matrix for NOEs

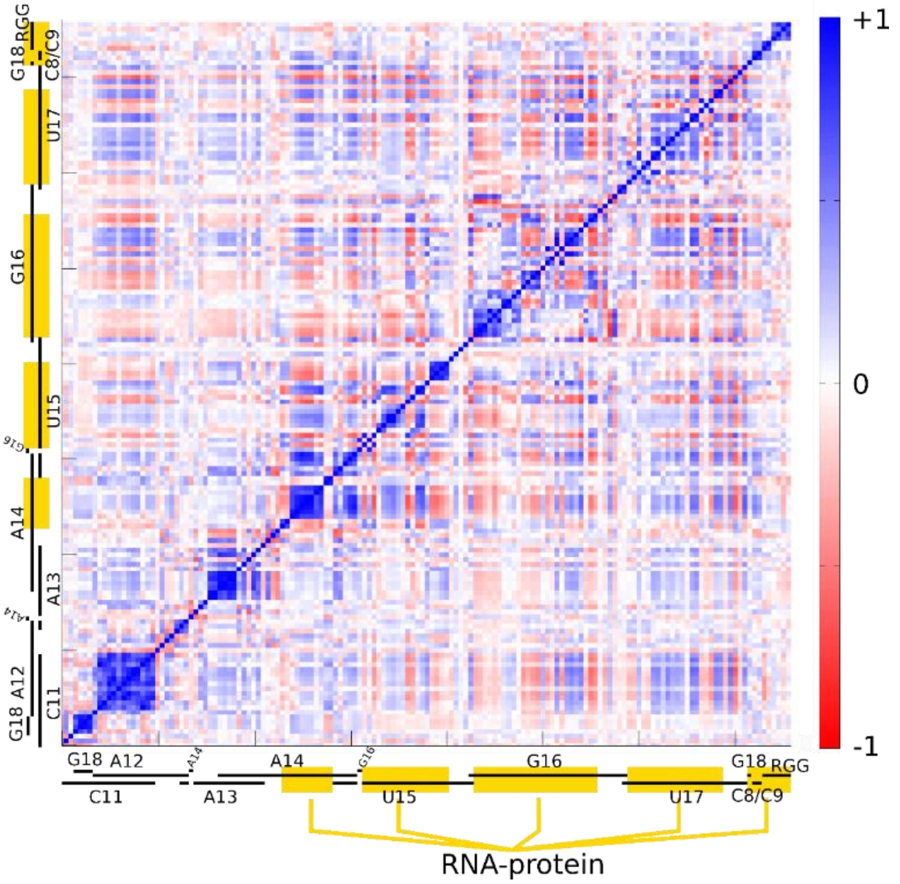

Figure S 8: Pearson correlation coefficients for $r^{-6}$ distances of experimentally measured NOE pairs over the whole FUS-U1 SL3 **S** ensemble for RNA loop region (RNA-RNA and RNA-protein NOEs). Black lines by the left and bottom edge show RNA residues to which the individual NOEs correspond while interactions with proteins are shown by yellow boxes; when only one partner is specified then it corresponds to intra-nucleotide NOEs. Spatially close regions with anti-correlations (mutually contradicting NOEs) suggest presence of multiple local sub-states. Improper sampling of these sub-states can lead to NOE violations in the ensemble if their populations are far from the real ones. Visible in the graph is also possible coupling along the RNA chain, i.e., the horizontal patterns of (anti)correlations.

### $U_{15}$-protein H-bond network instabilities in MD

For $U_{15}$ the simulations of both systems alternate experimentally-observed $U_{15}$(N3-H3)-Thr326(O) H-bond with $U_{15}$(N3-H3)-Thr326(OG). The latter H-bond is correlated with better H-bonding geometry of $U_{15}$(O2)-Thr326(N-H) H-bond in U1 SL3. However, states with $U_{15}$(N3-H3)-Thr326(OG) H-bond are coupled to $U_{15}$(H2′)-Phe288(HZ) NOE violations and are associated with $U_{15}$ movement towards $A_{14}$ in the pocket. This movement seems to be driven by $A_{14}$-$U_{15}$ stacking, which can be over-stabilized by the force field.

### Further analyses of RNA-RGG interactions

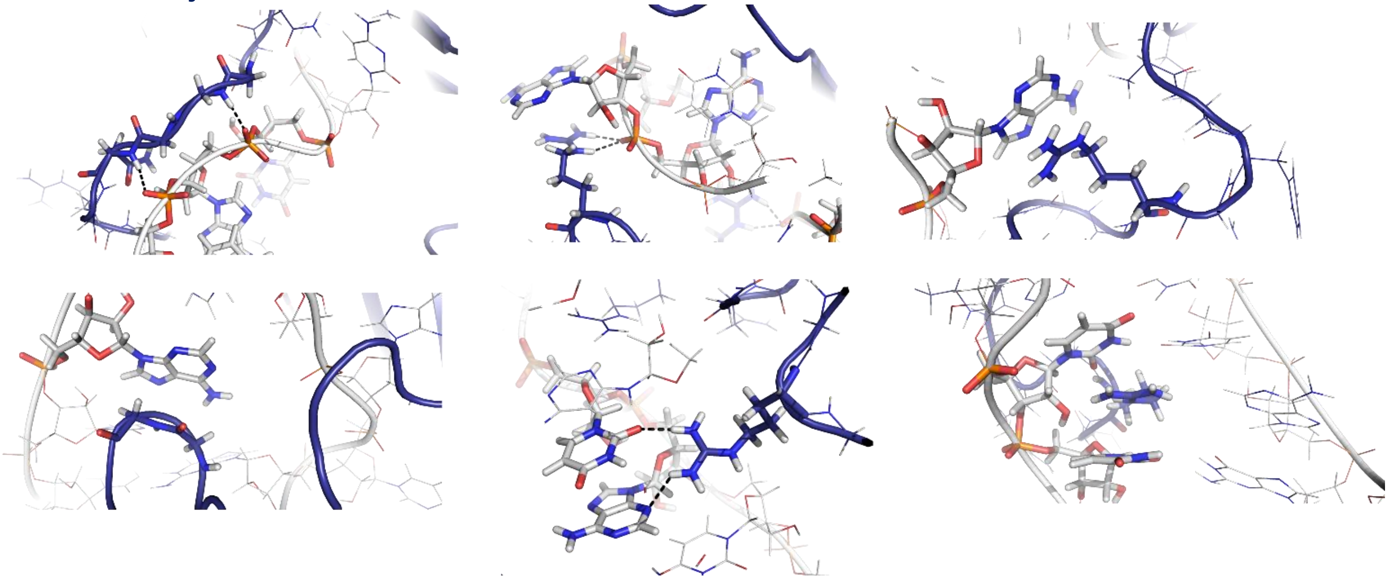

Figure S 9: Examples of some of the very many RGG-RNA interactions sampled in MD. Protein is in dark blue, RNA is white, H-bonds are shown as dashes. From upper-left to lower-right figure: H-bonding between RGG and RNA backbone; arginine-phospate salt-bridge combined with arginine-adenine stacking; arginine-adenine stacking; glycine-adenine stacking; simultaneous arginine H-bonding to two stacked bases; arginine intercalation in the hnRNPA2/B1 stem.

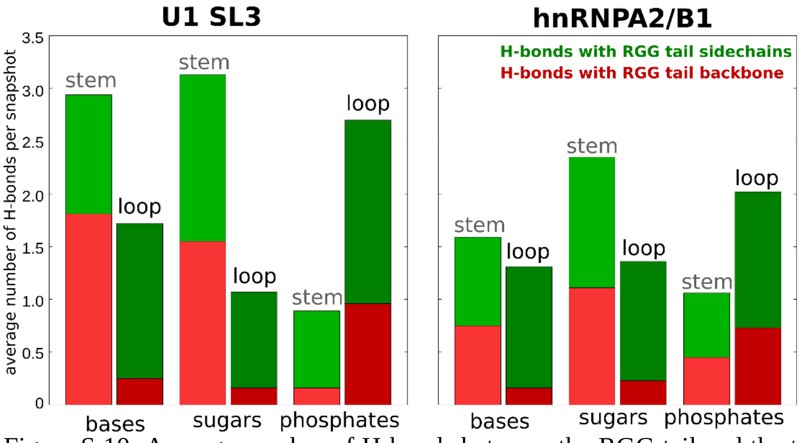

Figure S 10: Average number of H-bonds between the RGG tail and the two RNA hairpins. H-bond criterion was acceptor-donor distance < 3.5 Å and angle > 120˚. The graph shows that for U1 SL3 stem the RGG binds mainly to sugars and bases while for the loop region it binds rather to phosphates. This difference is less pronounced for the more flexible hnRNPA2/B1 hairpin. Interactions formed between RGG and RNA backbone are more frequent than for RNA bases. Note, that the balance of RGG-phosphate interactions is affected by the cufix correction compared to a standard force field, however, we suggest the cufix is more realistic.

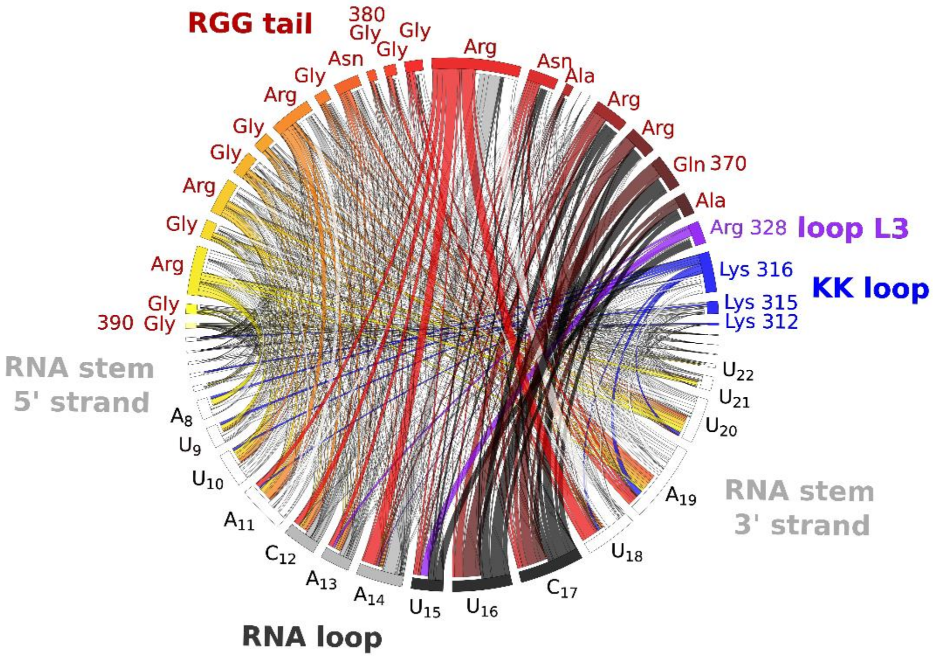

Figure S 11: Intermolecular H-bond network map for FUS-hnRNPA2/B1 complex; see the Main text Figure 9 for details. The only significant difference between RGG behavior between U1 SL3 and hnRNPA2/B1 complexes is the enhancement of Arg377 H-bonds to RNA in the latter (this arginine often enters the loop interior, see the "Excessive arginine stacking in FUS-U1 SL3 complex" section below).

### Analyses of stem flexibility and KK loop binding

Notable is the difference between flexibilities of the two RNA stems. The U1 SL3 stem has three GC WC base pairs just below the loop which are stable in all simulations. The hnRNPA2/B1 stem has a dynamic $U_{10}/A_{11}/U_{18}$ element below the loop followed by two AU WC base pairs and the GU wobble. The dynamics seen in this region of hnRNPA2/B1 includes reversible re-modelling of the base pairing up to GU wobble (see below) and enables protein to contact the RNA stem more extensively and variably than in the case of U1 SL3 (*Figure S 12*). Namely, we have observed reversible $A_{11}$ unpairing and switches from Watson–Crick $A_{11}$-$U_{18}$ to non-Watson–Crick $U_{10}$-$U_{18}$ base pairing (*Figure S 13*A and B). Further, U9-A19 pair undergoes reversible transitions to non-Watson–Crick base-pairing, with the non-Watson–Crick pairing being often supported by Lys36(O) H-bond to $U_9$(N3) or $A_{19}$(N6) (*Figure S 13*D). Similarly, GU wobble ($G_7$-$U_{21}$) frequently transits from conventional to bifurcated GU wobble, which is typically accompanied by Lys35(NZ) or RGG residues (Arg108/Gly109) H-bonding to the respective bases (*Figure S 13*C). Since all these altered RNA states were observed also in auxiliary simulations of free RNA, albeit with shorter lifetimes about singles of nanoseconds, we suggest that the protein binding only captures different available RNA conformations without inducing new ones.

We additionally tried to combine stems and loops of both systems which can be done in several ways as depicted in *Figure S 14*. When the hnRNPA2/B1 stem was swapped to the U1 SL3 stem while retaining loop size of seven nucleotides (the same size as in U1 SL3, *Figure S 14*D), we have observed increased propensity of $A_{13}$ bulging out compared to simulations of a similar construct but with only six nucleotides in the loop (the same size as the hnRNPA2/B1 loop, *Figure S 14*C). In general, it looks that a stiffer stem is in some conflict with the seven-nucleotide loop having all 5′-end bases inside, promoting thus bulging out of some of the loop nucleotides. In contrast, the more flexible stem absorbs the strain by its own deformation or plasticity.

The stem of the bound RNA is contacted and neutralized by RRM KK loop and RGG tail. The simulations show, that the KK-loop lysines can transiently contact the stem along its whole length. While we analyzed only direct H-bonds between KK loop lysines and RNA stem we also expect water-bridges to mediate the KK loop-RNA binding as water bridging is common for ion-pair interactions. As discussed in the Main text, the flexibility of U1 SL3 stem is increased by the $C_{24}$ bulge (illustrated by broader spread of stem bending angle in Figure S *15*A; helical axes were analyzed with x3DNA-DSSR[11]). Simulations of the isolated U1 SL3 stem with and without the $C_{24}$ bulge show that including the $C_{24}$ results in larger bending angle of the stem, bringing it closer to the KK loop (Figure S *15*). Further comparison of simulations of the isolated stems and of stem-loops bound to FUS shows that the ensemble of conformations (i.e., the stem bending) sampled by the stem is affected by the protein binding (Figure S *15*A).

We suggest, that KK loop lysines are important for correctly orienting the protein with respect to the RNA stem-loop, based on following observations. We performed 10x5 μs and 4x5 μs (for U1 SL3 and hnRNPA2/B1 FUS complexes, respectively) simulations with lysines of KK loop mutated to alanines. In four out of the ten FUS-U1 SL3 simulations the RRM-RNA binding interface was completely lost on time-scales ranging from 100 ns to 3 μs. In these cases the whole YNY motif unbound while in disruptions in Main text Figure 3 and SI *Table S 3* the loss of interface was mostly manifested by lossing one or two RRM-bound nucleotides. The losses of the whole YNY motif were accompanied by KK loop departures far away from the RNA stem and changes in the overall orientation of RNA towards RRM. In other two simulations, $U_{15}$ irreversibly unbound from its pocket. Binding interface in the FUS-hnRNPA2/B1 simulations with alanines in KK loop was still reproduced at the ends of the four simulations after 5 μs.

Importance of the KK loop-stem interactions in positioning the RNA is further visualized by simulations of short U1 SL3 ssRNA bound on the RRM. For these simulations, we have removed the RNA 5′-end of the RNA loop, RNA stem and RGG tail to decouple their effects from the YNY motif binding at the RNA-RRM binding interface. Set of 1-μs simulations shows that while binding of the A14-U16 on the interface was retained, $U_{17}$ quickly lost its position in the pocket. Only when we introduced a short double helix at the end of our ssRNA construct (3x CG base pair), the $U_{17}$ was stabilized in the pocket (Figure S 16).

In general, the simulations indicate long range communication between different parts of the FUS-RNA systems which is affected by internal flexibilities of the participating elements (this is visible mainly for the stems) and which si affected by the RNA sequence. For example, properties of the stem may affect the optimal length of the loop and the propensity of its bases to bulge out and search of their binding partners. It is even possible that when the bulge out bases interact with some another molecule, it could be sensed by the KK loop through its interaction with the stem.

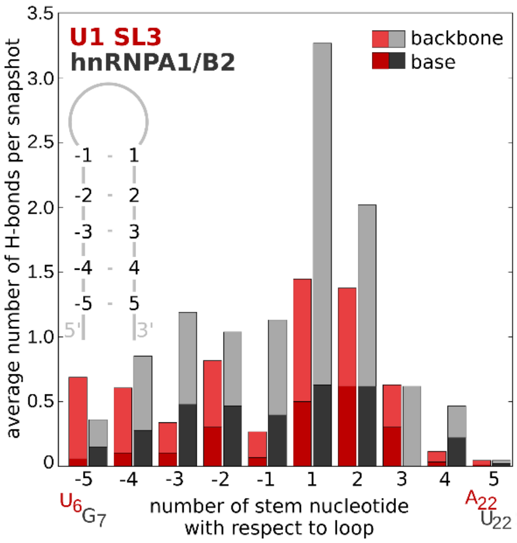

Figure S 12: Average number of H-bonds between RNA stem residues and FUS RRM (i.e., the RGG tail was excluded from the analysis) per snapshot. Criterion was donor-acceptor distance < 3.5 Å. Average numbers can exceed 1 since multiple different H-bonds between nucleotide and RRM can be present in one snapshot.

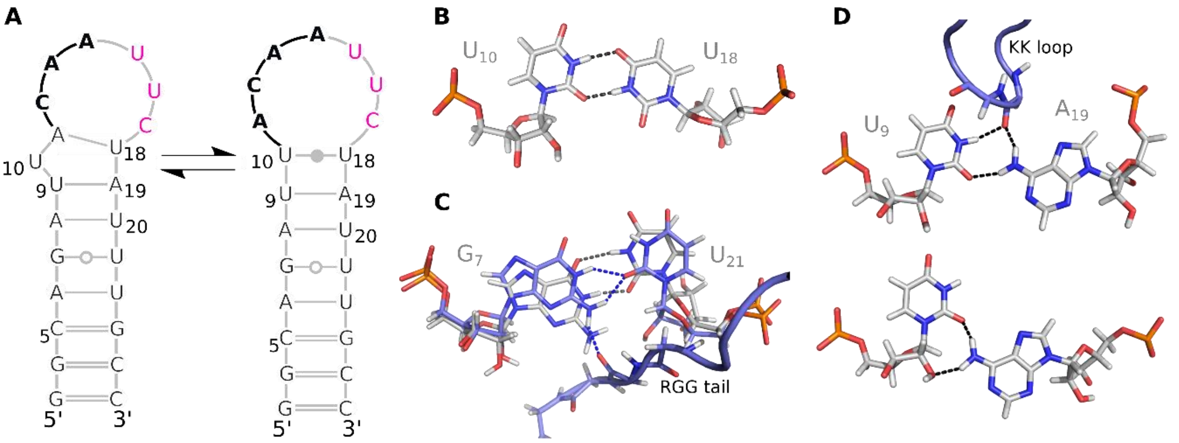

Figure S 13: Examples of hnRNPA2/B1 dynamics (substates) in the simulations. **A** Formation of U10-U18 base pair. **B** U10-U18 base pair structure. Sometimes, RGG tail arginines H-bonded to major groove donors. **C** G7-U21 wobble geometries: conventional geometry (gray) and bifurcated one (blue), which is stabilized by the RGG tail. **D** two cases of U9-A19 non-Watson–Crick base pairing.

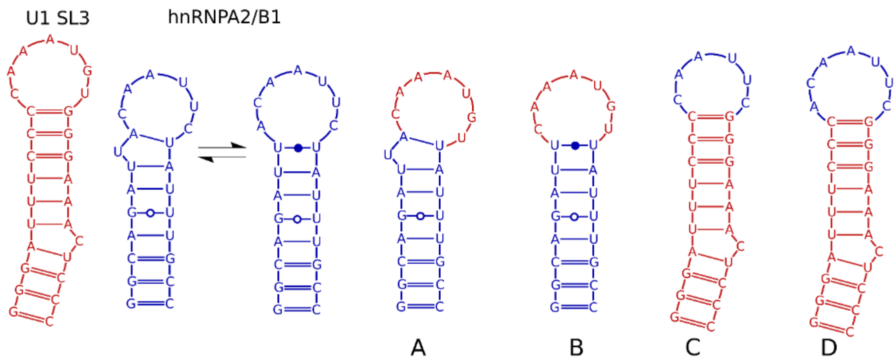

Figure S 14: Tested constructs (A-D) of the stem-swapping.

Table S 6: Simulation behavior (A13 bulging out and interface disruptions) in stem-swapped simulations.

| MD (5 µs) | **system C** - hnRNPA2/B1 loop with U1 SL3 stem, 6 nucleotide loop ($C_{12}$-$C_{17}$) | **system D** - hnRNPA2/B1 loop with U1 SL3 stem, 7 nucleotide loop ($A_{11}$-$C_{17}$) |
|---|---|---|
| | simulation behavior | |
| 1 | - | $A_{13}$ bulged out at 20 ns |
| 2 | disruption of RNA-RRM interface ($U_{16}$) at 700 ns | $A_{13}$ bulged out at 40 ns |
| 3 | - | $A_{13}$ bulged out at ~3000 ns |
| 4 | $A_{13}$ bulged out at ~2000 ns | disruption of RNA-RRM interface (stem-$U_{17}$ binding) at 100 ns |
| 5 | $A_{13}$ bulged out at ~1000 ns, bulged back in at ~4000 ns | $A_{13}$ bulged out at 800 ns |
| 6 | disruption of RNA-RRM interface ($U_{16}$) at ~3400 ns | disruption of RNA-RRM interface ($U_{16}$) at 150 ns |
| 7 | $A_{13}$ bulged out at ~3500 ns | $A_{13}$ bulged out at 250 ns, disruption of RNA-RRM interface ($U_{17}$) at ~1500 ns |

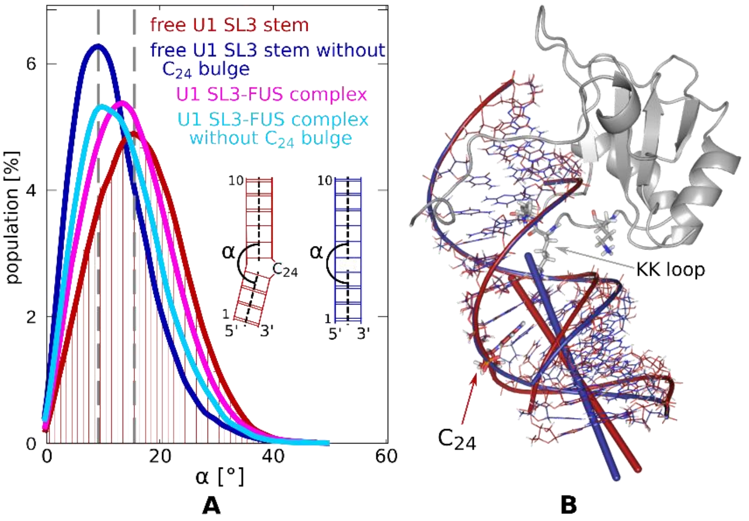

Figure S 15: U1 SL3 stem flexibility. **A** Distribution of angle alpha between two helical axes (defined in the figure) in simulations of model stems and in complexes with protein with and without $C_{24}$ bulge. The bin size for creating the distribution curves was 1˚ (bins are shown for the free U1 SL3 stem). The gray vertical dashed lines mark the maximums for simulations of free stems. **B** Comparison of average structures from free stem simulations of U1 SL3 with and without $C_{24}$ bulge. The structures were aligned with respect to the upper stem part (six base pairs above the bulge) and the bars show helical axes of lower helices (see **A** for definition of axes). Protein position is also shown for orientation though the protein was not included in these free stem simulations; KK loop lysines are shown as sticks.

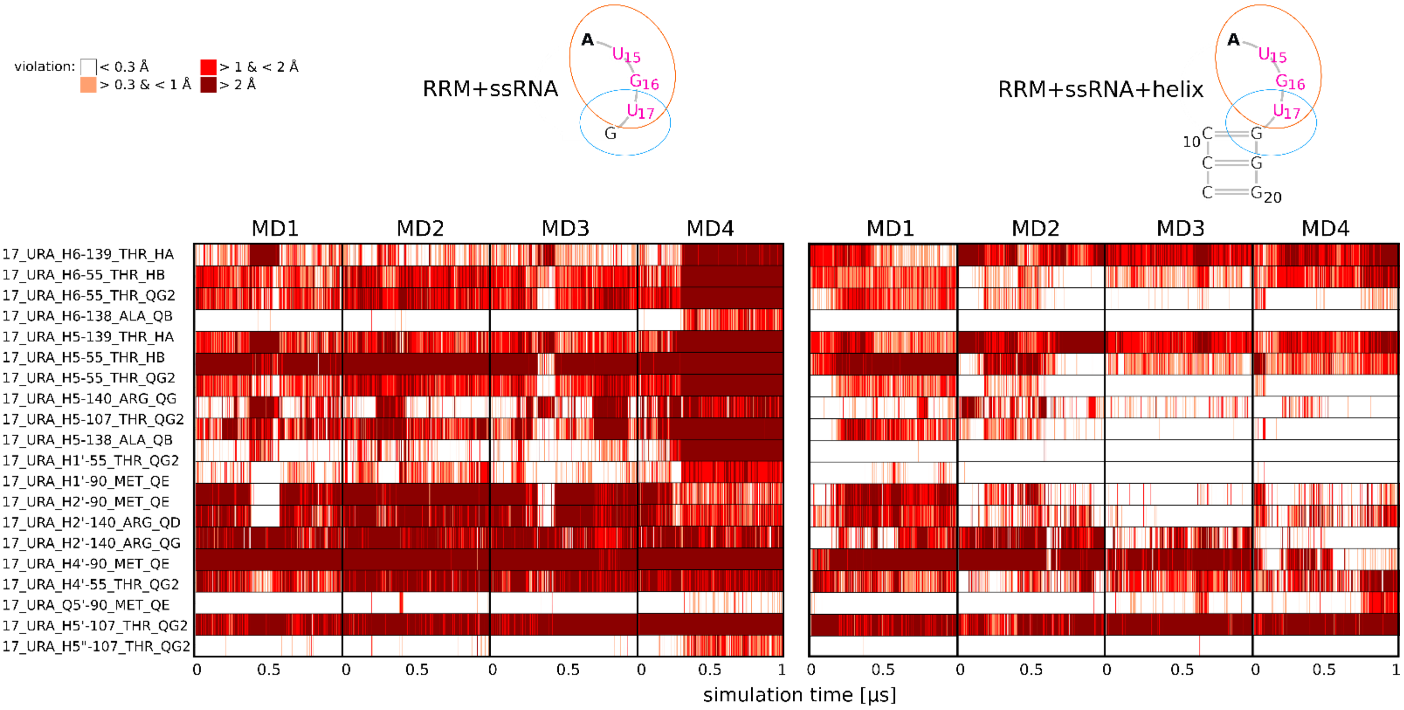

Figure S 16: $U_{17}$ binding in simulations with two different short RNA constructs. The RNA constructs are show on the schemes above graphs. $U_{17}$ position in RRM pocket is visualized by monitoring instantaneous RNA-protein NOE violations during the courses of four independent trajectories.

### Excessive arginine stacking in the FUS-U1 SL3 complex

*Preliminary* simulations of the FUS-U1 SL3 complex revealed frequent formation of stable arginine Arg383 sidechain stacking with $G_{18}$, accompanied by $C_{11}$-Arg383 H-bond pairing (*Figure S 17*A). In majority of cases the $C_{11}$-Arg383 pairing kinetically preceded the stacking. This interaction effectively extends the RNA stem and prevents $C_{11}$ from bulging out to the solvent. Once formed, it remained stable in all our trajectories (irreversible on 5-μs time scale in multiple trajectories) and it thus became the dominant Arg383 state. This led to restriction of RGG tail dynamics (*Figure S 17*B). Such arginine pseudo-base-pairing interaction is an analogue to what is seen in some DNA-protein repair complexes. However, in our case the conformation violates several NOE signals (some $C_{11}$-$A_{12}$, all $C_{11}$-$G_{18}$, and some RGG tail-RNA signals). Thus, despite that formation of this arginine stacking may be realistic, its large population would be inconsistent with the experimental data. Minor population of this interaction, however, cannot be ruled out.

Nevertheless, it is evident that the Arg383-$C_{11}$/$G_{18}$ interaction is over-stabilized by the force field and thus we included all trajectory portions where this interaction is present to the **U** ensemble.

Thus, in order to eliminate this spurious stacking we carried out all simulations in the *main* set of simulations with enhanced Arg383-water interactions using the NBfix approach by scaling $R_{min}$ for N-$O_w$ atom pairs by 0.95 (see Methods). Formation of Arg383 stacking was almost eliminated, except for occasional short transient events. Our destabilization of Arg383 may be excessive but we intended to have more sampling of structures without this stack rather than fine-tuning/balancing the stacking.

In case of the FUS-hnRNPA2/B1 complex, no arginine formed an analogous permanent stacking interaction. This may be due to the fact, that hnRNAPA1/B2 hairpin does not possess the stiff segment with consecutive three GC base pairs producing the stackign platform in the stem as its stem is more flexible. Indeed, when we swapped the hnRNPA2/B1 stem to U1 SL3 stem (i.e. $U_9$-$A_{11}$+$U_{18}$-$A_{19}$ part became a CG stem, *Figure S 14*C), we observed in four out of seven simulations the same arginine-stacking interaction as described above, albeit the stacked arginine was Arg377 rather than Arg383. Arg377 was observed to go inside the hnRNAPA1/B2 loop also in simulations with the native hairpin, where it reversibly stacked mainly with U18 but also with A11.

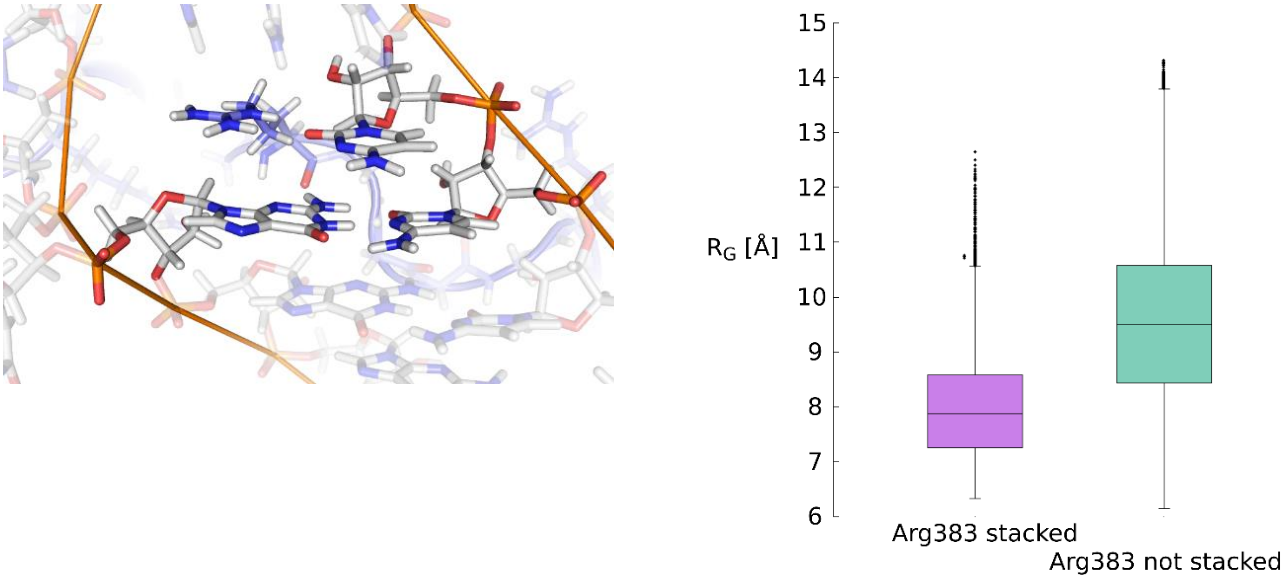

**A** **B**

Figure S 17: **A** Arg383 forming pseudopair with $C_{11}$ and stacking on top of the $C_{10}$-$G_{18}$ base pair. **B** Radii of gyration ($R_G$) of RGG tail in trajectory parts with Arg383 stacked and not stacked on top of the $C_{10}$-$G_{18}$ pair.
